# Explaining decisions of Graph Convolutional Neural Networks: patient-specific molecular subnetworks responsible for metastasis prediction in breast cancer

**DOI:** 10.1101/2020.08.05.238519

**Authors:** Hryhorii Chereda, Annalen Bleckmann, Kerstin Menck, Júlia Perera-Bel, Philip Stegmaier, Florian Auer, Frank Kramer, Andreas Leha, Tim Beißbarth

**Affiliations:** Medical Bioinformatics, University Medical Center Göttingen, Germany; Dept. of Medicine A (Hematology, Oncology, Hemostaseology and Pulmonology), University Hospital Muenster, Germany; Hospital del Mar Medical Research Institute (IMIM) Barcelona, Spain; geneXplain GmbH, Wolfenbüttel, Germany; IT Infrastructure for Translational Medical Research, University of Augsburg, Germany; Medical Statistics, University Medical Center Göttingen, Germany

**Keywords:** gene expression data, explainable AI, personalized medicine, precision medicine, classification of cancer, deep learning, prior knowledge, molecular networks

## Abstract

**Motivation:** Contemporary deep learning approaches show cutting-edge performance in a variety of complex prediction tasks. Nonetheless, the application of deep learning in healthcare remains limited since deep learning methods are often considered as non-interpretable black-box models. Layer-wise Relevance Propagation (LRP) is a technique to explain decisions of deep learning methods. It is widely used to interpret Convolutional Neural Networks (CNNs) applied on image data. Recently, CNNs started to extend towards non-euclidean domains like graphs. Molecular networks are commonly represented as graphs detailing interactions between molecules. Gene expression data can be assigned to the vertices of these graphs. In other words, gene expression data can be structured by utilizing molecular network information as prior knowledge. Graph-CNNs can be applied to structured gene expression data, for example, to predict metastatic events in breast cancer. Therefore, there is a need for explanations showing which part of a molecular network is relevant for predicting an event, e.g. distant metastasis in cancer, for each individual patient.

**Results:** We extended the procedure of LRP to make it available for Graph-CNN and tested its applicability on a large breast cancer dataset. We present Graph Layer-wise Relevance Propagation (GLRP) as a new method to explain the decisions made by Graph-CNNs. We demonstrate a sanity check of the developed GLRP on a hand-written digits dataset, and then applied the method on gene expression data. We show that GLRP provides patient-specific molecular subnetworks that largely agree with clinical knowledge and identify common as well as novel, and potentially druggable, drivers of tumor progression. As a result this method could be potentially highly useful on interpreting classification results on the individual patient level, as for example in precision medicine approaches or a molecular tumor board.

**Availability:** https://gitlab.gwdg.de/UKEBpublic/graph-lrp https://frankkramer-lab.github.io/MetaRelSubNetVis/

## 1 Introduction

Gene-expression profiling as for example by DNA microarrays or next generation sequencing are becoming more and more available as the technologies become cheaper and quicker. As a result, high-throughput technologies played a significant role in identifying predictive gene signatures and discovering individual biomarkers in cancer prognosis (Perera, Leha, and Beissbarth, 2019) Furthermore, high-throughput sequencing produces huge amounts of data that can be used for deriving clinical predictors for relapse events (e.g occurrence of metastases). At the moment, deep learning techniques have shown prominent results in many research fields with big and complex data.

In recent years deep learning was applied to a wide range of problems in various areas. Deep learning is a class of machine learning methods based on neural networks that are aimed at the automatic learning of data representations (features) from raw data. These neural network methods demonstrated state-of-the-art performance in visual object recognition, object detection and speech recognition. The deep learning breakthroughs were made mostly on the data that have underlying Euclidean structure. One of the successfully used deep learning methods applied on image data are convolutional neural networks (CNNs) that exploit the grid-like structure of images. In many cases data is structured in non-Euclidean domains as well, for example networks in social sciences and molecular networks in biology. Recently, deep learning methods extended to domains like graphs and manifolds (Monti et al., 2017) Conventional deep learning and CNNs are already used in the field of bioinformatics (Min, Lee, and Yoon, 2016) drug discovery and genomics (LeCun, Bengio, and Hinton, 2015) Deep learning on graphs inspired further developments showing promising results on metastatic events prediction (Chereda et al., 2019) and subtypes classification (Rhee, Seo, and Kim, 2018) in breast cancer, on modeling drug-drug interactions (Ma et al., 2018; Zitnik, Agrawal, and Leskovec, 2018) and predicting protein-protein interactions (Leskovec, 2018)

Remarkably, deep neural networks are able to model complex interaction between the input and output variables. Furthermore, multiple hidden layers create multiple interactions between input features. This complexity does not allow easy tracking of how a fixed input feature influences the output, thus a neural network itself as a black-box machine learning model does not give interpretable insights.

Decisions proposed by neural networks have to be explained for the application in the clinical domain (Yang et al., 2018) Furthermore, the European Union’s new General Data Protection Regulation (GDPR) restricted automated decision making produced by, e.g., algorithms (*2018 reform of EU data protection rules* 2018) Article 13 *Information to be provided where personal data are collected from the data subject* specifies that the data controller (e.g. clinics) should provide the data subject (e.g. patients) with “meaningful information about the logic involved”. Article 22 *Automated individual decision-making, including profiling* states that “The data subject shall have the right not to be subject to a decision based solely on automated processing”, unless the data subject gives a consent with it (paragraph 2.c). Therefore, the combination of explainability and the expressiveness of deep neural networks is yet a task to work on (Yang et al., 2018)

Explanation methods for complex nonlinear models such as neural networks (including convolutional) aim to interpret classification decisions of a machine learning model in terms of input variables. These methods can be categorized into two groups (Montavon et al., 2017) functional approaches and message passing approaches. The first group of methods produce explanations out of local analysis of a prediction including the sensitivity analysis, Taylor series expansion, and model agnostic approaches LIME (Ribeiro, Singh, and Guestrin, 2016) and SHAP (Lundberg and Lee, 2017) The second group provides explanations by running a backward pass in a computational graph, which generates a prediction as its output. The Layer-Wise Relevance Propagation (LRP) method (Bach et al., 2015) combines functional and message passing approaches to generate relevances of each input feature. For a fixed input feature, the relevance shows how much this feature influences the classifier decision. The relevances are generated for each data point individually, which is a huge advantage of this explainability method.

One of the tasks of clinical cancer research is to identify prognostic gene signatures that are able to predict clinical outcome (Johannes et al., 2010) From a machine learning perspective, the endpoint is usually presented as a classification task, and the challenge is to find discriminative features. However, the search for such molecular markers is based on high-dimensional datasets, where the number of genes is much higher than the number of patients. The “curse of dimensionality” leads to instability in the feature selection process. Improvements can be made by including prior knowledge of molecular networks (e.g. pathways) into a machine learning algorithm. According to (Johannes et al., 2010) the machine learning methods benefit from pathway knowledge since genes are not treated as independent. This benefit is based on the hypothesis that neighboring genes should have similar expression profiles. Consequently, the decision of machine learning methods is formed by predictive subnetworks. Furthermore, these subnetworks can differ from one patient to another according to their expression profiles. Convenient feature selection methods, that are utilizing prior knowledge (Johannes et al., 2010; Binder and Schumacher, 2009) provide general features that are the same for all patients. However, we adapted an existing LRP technique to Graph-CNN (Defferrard, Bresson, and Vandergheynst, 2016) which can incorporate a molecular network. Thus, we provide patient specific subnetworks that are individual for each patient. According to the knowledge of the authors, a feature selection method that benefits from prior knowledge and provides patient-specific subnetworks has not been shown before.

There are some recent interpretation methods specialized for graph-neural networks. (Xie and Lu, 2019) and (Pope et al., 2019) provide explainability methods that are exactly based on and crafted only for Graph Convolutional Network of (Kipf and Welling, 2016) utilizing convolutional architecture which is a simplified version of that of Graph-CNN (Defferrard, Bresson, and Vandergheynst, 2016) we use. Ying et al suggested a model-agnostic GNNExplainer that is suitable for node classification, link prediction, and graph classification. The essence of our classification task is to predict an occurrence of distant metastasis based on gene expression data structured by a protein-protein interaction network. Since each vertex of a molecular network has a corresponding gene expression value as an attribute, we perform a graph-signal classification task. Ying et al demonstrates a solution to explaining node classification, but do not provide an application of their approach for a graph-signal classification task. (Chereda et al., 2019) and (Rhee, Seo, and Kim, 2018) already applied Graph-CNN (Defferrard, Bresson, and Vandergheynst, 2016) in this context. Hence, there is still a lack of methods explaining Graph-CNN performing aforementioned machine learning task.

The novelty of our work consists of 2 parts. First, we present a method delivering data-point (i.e. patient) specific explanations for Graph-CNN (Defferrard, Bresson, and Vandergheynst, 2016) in the context of graph-signal classification. Second, we show how these patient-specific molecular subnetworks assist the need in personalized precision medicine decisions via explaining patient-specific predictions of Graph-CNN applied on a large breast cancer dataset. Breast cancer is the second most common cancer in industrialized countries (Bray et al., 2018) Patients often develop distant metastases that limit survival due to the lack of curative treatment options (Bray et al., 2018) We interpret the classifier’s inferences by patient-specific subnetworks that would explain the differential clinical outcome and identify therapeutic vulnerabilities.

## 2 Materials and Methods

### 2.1 Breast Cancer Data

We applied our methods to a large breast cancer patient dataset that we previously studied and preprocessed (Bayerlová et al., 2017) That data is compiled out of 10 public microarray datasets measured on Affymetrix Human Genome HG-U133 Plus 2.0 and HG-U133A arrays. The datasets are available from the Gene Expression Omnibus (GEO) (Barrett et al., 2013) data repository and have the accession numbers GSE25066, GSE20685, GSE19615, GSE17907, GSE16446, GSE17705, GSE2603, GSE11121, GSE7390, GSE6532. The RMA probe-summary algorithm (Irizarry et al., 2003) was used to process each of the datasets, and only samples with metastasis-free survival were selected and combined together on the basis of HG-U133A array probe names. Quantile normalization was applied over all datasets. In the case of few probes mapping to one gene, the probe with the highest average value was taken. In the end, we ended up with 12179 genes per each patient. To formulate two classes for the prediction task we selected 393 patients with distant metastasis occurred within the first 5 years, and 576 patients without metastasis having the last follow up between 5 and 10 years. Breast cancer molecular subtypes for the patient samples were predicted in (Bayerlová et al., 2017) utilizing *genefu* R-package (Gendoo et al., 2016)

### 2.2 Protein-Protein Interaction Network

We used the Human Protein Reference Database (HPRD) protein-protein interaction (PPI) network (Keshava Prasad et al., 2009) to structure the gene expression data. The database contains protein-protein interaction information based on yeast two-hybrid analysis, in vitro and in vivo methods. The PPI network is an undirected graph with binary interactions between pairs of proteins. The graph is not connected. We mapped the genes from the gene expression data to the vertices of the PPI network. Resulting PPI graph has 7168 vertices (genes) matched, and 207 connected components. The main connected component has 6888 vertices, and each of the other 206 components has from 1 to 4 vertices. Further, we utilized only the main connected component since the Graph-CNN requires graphs to be connected.

### 2.3 Pathway Analysis

Enrichment of signal transduction pathways annotated in the TRANSPATH^®^ database version 2020.1 (Krull et al., 2003) in genes prioritized by GLRP were analyzed using the geneXplain platform version 5.1 (Koschmann et al., 2015) The analysis based on the Fisher’s exact test (Fisher, 1922) was carried out for gene sets obtained for individual patients as well as for their combination into subtype gene sets. The following calculations were applied to investigate differences in pathway hits. Let *P* denote a set of pathway genes and *S*_*i*_ and *S*_*k*_ two subnetwork gene sets, so that *P*_*i*_ = *P* ∩ *S*_*i*_ and *P*_*k*_ = *P* ∩ *S*_*k*_ are the sets of pathway genes matched by the two subnetworks. The difference ∩*P*_*i,k*_ in matched pathway genes was then calculated as |(*P*_*i*_ ∪ *P*_*k*_) *\* (*P*_*i*_ ∩*P*_*k*_)|*/*|*P*_*i*_ ∪ *P*_*k*_| with |*P*_*i*_ ∪ *P*_*k*_| *>* 0. For each selected pathway, we calculated Δ*P*_*i*_, *k* for each pair of subnetworks and reported the median of examined pairs.

### 2.4 Problem formulation

We focus on explaining classifier decisions of Graph-CNN adapting existing LRP approaches for graph convolutional layers. LRP should be applied as a postprocessing step to a model already trained for the machine learning task. The task is formulated as a binary classification of gene expression data *X* ∈ *R*^*n*×*m*^ to target variable *Y* ∈ {0, 1}^*n*^ representing the appearance of a distant metastatic event. *n* is a number of samples (patients) and *m* is a number of features (genes). The information of the molecular network is presented as an undirected graph *G* = (*V, E, A*), where *V* and *E* denote the sets of vertices and edges respectively. *A* is the adjacency matrix of dimensionality *m* × *m*. A row *x* of the gene expression matrix *X* contains data from one patient and can be mapped to the vertices of the graph *G*. In such a way, values of *x* are interpreted as a graph signal.

A trained neural network can be represented as a function 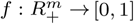 mapping the positive input to the probability of the output class. The input *x* is a set of gene expression values *x* = {*x*_*g*_ } where *g* denotes a particular gene. The function *f* (*x*) computes the probability that a certain pattern of gene expression values is present w.r.t to the output class. LRP methods apply propagation rules from the output of the neural network to the input in order to quantify the relevance score *R*_*g*_ (*x*) for each gene *g*. These relevances show how much gene *g* influences the prediction *f* (*x*) :

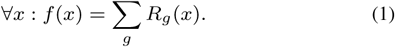

Equation (1) (Montavon et al., 2017) demonstrates that the relevance scores are calculated w.r.t every input data point *x*.

### 2.5 Graph Convolutional Neural Network and Layer-wise Relevance propagation

Usual CNNs learn data representations on grid-like structures. The Graph-CNN (Defferrard, Bresson, and Vandergheynst, 2016) as a deep learning technique is designed to learn features on graphs. The convolution on graphs is used to capture localized patterns of a graph signal. This operation is based on spectral graph theory. The main operator to investigate the spectrum of a graph is the graph Laplacian *L* = *D* − *A*, where *D* is a weighted degree matrix, and *A* is a weighted adjacency matrix. *L* is a real symmetric positive semidefinite matrix that can be diagonalized such that *L* = *U* Λ*U*^*T*^, where Λ = *diag* ([*λ*_1_, *…, λ*_*m*_]) is a diagonal non-negative real valued matrix of eigenvalues, matrix *U* is composed of eigenvectors. Matrices *U* and *U*^*T*^ define the Fourier and the inverse Fourier transform respectively. According to the convolution theorem, the operation of graph convolution can be viewed as a filtering operation:

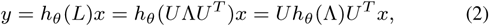

where *x, y* ∈ *R*^*m*^, and the filter *h*_*θ*_ (Λ) is a function of eigenvalues (graph frequencies). To localize filters in space, the authors in (Defferrard, Bresson, and Vandergheynst, 2016) decided to use a polynomial parametrization

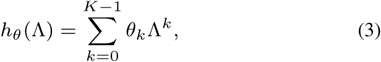

where *θ* ∈ *R*^*k*^ is a vector of parameters. The order of the polynomial, which is equal to *K* − 1, specifies the local *K* − 1 hop neighborhood. The neighborhood is determined by the shortest path distance. The polynomial filter can be computed recursively, as a Chebyshev expansion, which is commonly used in graph signal processing to approximate kernels (Hammond, Vandergheynst, and Gribonval, 2011) The Chebyshev polynomial *T*_*k*_ (*x*) of order *k* is calculated as *T*_*k*_ (*x*) = 2*xT*_*k*− 1_ (*x*) − *T*_*k*−2_(*x*) with *T*_0_ = 1 and *T*_1_ = *x*. The Chebyshev expansion applies for values that lie in [−1, 1], therefore, the diagonal matrix of eigenvalues Λ has to be derived from a rescaled Laplacian *L* = (*D* − *A*)*/λ*_*max*_ − *I*_*n*_. Thus, the filtering operation can be rewritten as

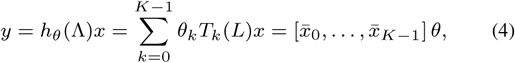

where 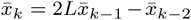 with 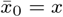 and *x*_1_ = *Lx*. The transition in equation 4 is done according to the observation (*U* Λ*U*^*T*^) ^*k*^ = *U* Λ^*k*^*U*^*T*^. The filtering at the convolutional layer boils down to an efficient sequence of *K* −1 sparse matrix-vector multiplications and one dense matrix-vector multiplication (Defferrard, Bresson, and Vandergheynst, 2016)

LRP is based on the theoretical framework of deep Taylor decomposition. The function *f* (*x*) from equation (1) can be decomposed in terms of the Taylor expansion at some chosen root point *x*^∗^ so that *f* (*x*^∗^) = 0. The first order Taylor expansion of f(x) is:

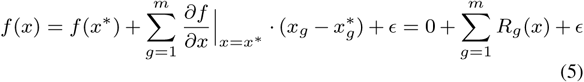

where the relevances *R*_*g*_ (*x*) are the partial differentials of the function *f* (*x*). The details of how to choose a good root point are described in Montavon et al., 2017. The *f* (*x*) represents a neural network which consists of multiple layers and each layer consists of several neurons. A neuron receives a weighted sum of its inputs and applies a nonlinear activation function. The idea of the deep Taylor decomposition is to perform a first order Taylor expansion at each neuron of the neural network. These expansions allow to produce relevance propagation rules that compute relevances at each layer in a backward pass. The rules redistribute the relevance from layer to layer starting from output until the input is reached. The value of the output represents the model’s decision which is equal to the total relevance detected by the model.

LRP is commonly applied to deep neural networks consisting of layers with rectified linear units (ReLU) nonlinearities. In our experiments, we use only this activation function. Let *i* and *j* be the neurons at two consecutive layers at which the relevance should be propagated from *j* to *i*. The neurons are of the type:

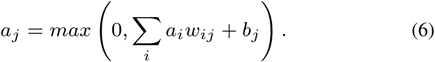

Noticeably, the layers of this type always have non-negative activations. The relevance propagation rule is the following:

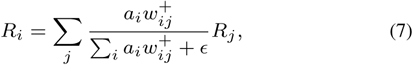

where 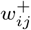 corresponds to the positive weights *w*_*ij*_ and *ϵ* stabilizes numerical computations (Yang et al., 2018) We set *E* to 1^−10^. Equation (7) depicts the *z*^+^ rule coming from deep Taylor decomposition (Montavon et al., 2017) The *z*^+^ rule is commonly applied to the convolutional and fully connected layers. It favours the effect of only positive contributions to the model decisions. The first input layer can have other propagation rules that are specific to the domain (Montavon, Samek, and Müller, 2018) In our work we used the rule (7) for the input layer as well since the gene expression data has positive values.

In order to propagate relevance through the filtering (4) we rewrite it as follows:

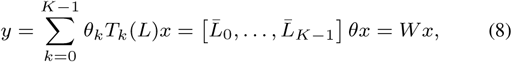

where matrix *W* ∈ *R*^*m*×*m*^ connects nodes *y* and *x*. The computation of matrix *W* is done as: 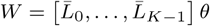, where 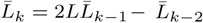 with 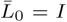 and 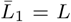 are the Chebyshev polynomials of the Laplacian matrix.

Each convolutional layer has *F*_*in*_ channels 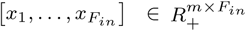 in the input feature map and *F*_*out*_ channels [*y*_1_, *…, yF*_*out*_] ∈ *R*^*m*^ × *F*_*out*_ of the output feature map. We consider the values of output feature maps before applying ReLU non-linearities on them. The *F*_*in*_ × *F*_*out*_ vectors of the Chebyshev coefficients *θ*_*i,j*_ ∈ *R*^*k*^ are the layer’s trainable parameters. The input feature map can be transformed into a vector 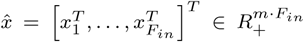. We adapt equation (8) to compute the *j*^*th*^ channel of the output feature map based on the input feature map:

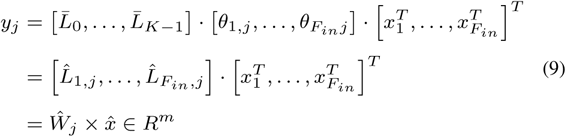

where 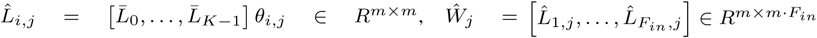

Since the *j*^*th*^ channel of the output feature map is connected through the matrix-vector multiplication with the input feature map, *Ŵ* _*j*_ can be treated as a matrix of weights joining two fully-connected layers. Therefore, the relevance 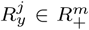 from the *j*^*th*^ output channel can be propagated to the input feature map relevance 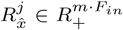 according to the rule (7). Overall, the relevance propagated from the output feature map to the input feature map is:

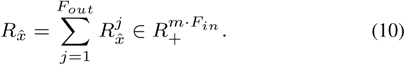

For running LRP on graph convolutional layers one needs to compute huge and dense matrices 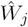. It requires *K* − 2 sparse matrix-matrix multiplications and one sparse to dense matrix-matrix multiplication. The computations for relevance propagation are heavier and much more memory demanding compared to the filtering (4).

### 2.6 Validation

To check the biological relevance of subnetwork genes prioritized by GLRP we also used the gene expression data from human umbilical vein endothelial cells (HUVECs) treated or not treated with tumor necrosis factor alpha TNFα (Rhead et al., 2020) The data, provided by the same authors (GEO database series: GSE144803), is suitable for a binary classification task and is balanced. The data was quantile normalized and mapped to the vertices of HPRD PPI resulting in 7798 genes in the main connected component. We compared gene sets identified in our subnetworks to gene modules and differentially expressed genes in response to TNFα identified by (Rhead et al., 2020) (Supplementary File S1). (Rhead et al., 2020) used weighted gene co-expression network analysis (WGCNA) (Zhang and Horvath, 2005) constructing networks as gene modules. Associations between subnetwork genes sets and 16 gene modules defined by (Rhead et al., 2020) as well as 589 upregulated genes (log-fold change > 0.5, FDR < 0.01), 425 downregulated genes (log-fold change < -0.5, FDR < 0.01) and the combined set of 1014 DE genes were analyzed using the *Functional classification* tool of the geneXplain platform (Kolpakov et al., 2011) Fisher test calculations were carried out with a total contingency table count corresponding to the number of genes in (Rhead et al., 2020, file S1 of) after mapping to Ensembl (Yates et al., 2020) gene ids (10022 genes).

## 3 Results

### 3.1 Sanity check of the implemented graph LRP

To initially validate our implemented LRP we applied Graph-CNN on the MNIST dataset (Lecun et al., 1998) in the same way as described in the paper (Defferrard, Bresson, and Vandergheynst, 2016) The MNIST dataset contains 70,000 images of hand-written digits each having a size of 28 by 28 pixels. To apply Graph-CNN on the image data, we constructed an 8 nearest-neighbors graph similarly to the schema proposed in (Defferrard, Bresson, and Vandergheynst, 2016) with the exception that all the weights are equal to 1. The weight 1 is more natural for the graph connecting neighboring image pixels. Thus, each image is a graph signal represented by node attributes - pixel values. We achieved high classification accuracy (99.02%) on the test set for the Graph-CNN, which is comparable to the performance of classical CNN (99.33%) reported in (Defferrard, Bresson, and Vandergheynst, 2016) The number of parameters were the same for both methods.

Usually, to manage box-constrained pixel values, the special pixel specific LRP rule is applied for the input layer (Montavon et al., 2019) This pixel specific rule highlights non only the digits itself, but also the contours of the digits (Montavon, Samek, and Müller, 2018, Figure 13 of) In contrast, the rule (7) highlights only those positively relevant parts of the image where the signal of the digit is present. We kept the propagation rule (7) for the input and all other layers in all our experiments. Further, we visually compared on the same digits how the heatmaps generated by implemented GLRP correspond to the heatmaps generated by usual LRP procedure applied on classical CNN (figure 1).

**Fig. 1.**
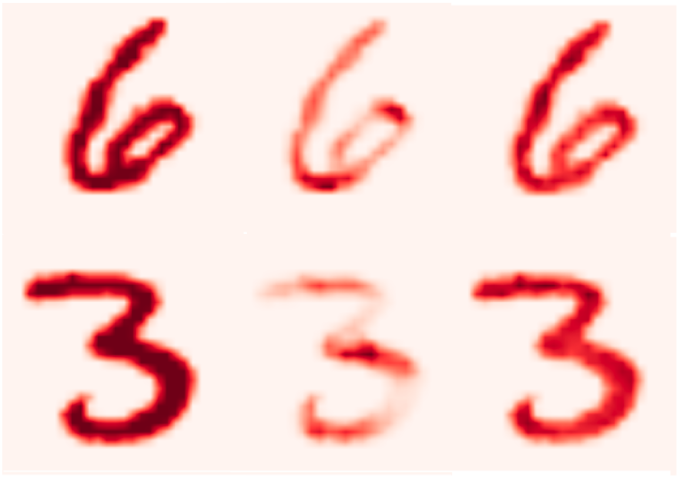
From left to right: initial image, LRP on classical CNN and GLRP on Graph-CNN.

The heatmaps were rendered only for the classes predicted by classical CNN and Graph-CNN. In this case the classes are “6” and “3”. For the Graph-CNN a bigger part of the digit is relevant for the classification since the covered neighborhood can be expanded up to 24 hops. Graph-CNN’s filters are isotropic, thus they tend to cover roundish areas that concern rounded patterns (curves) of the digit (Supplementary Figure S1).

### 3.2 Genes selected by GLRP correlate with modules identified by gene co-expression network analysis

We trained a Graph-CNN on gene expression data mentioned in the section 2.6 to classify HUVECs treated or not treated with TNFα. We utilized Graph-CNN architecture consisting of 2 convolutional layers with 4 and 8 filters respectively followed by one hidden fully connected layer with 128 nodes. No pooling was used. The performance of the method is the same as for Random Forest in 10-fold cross validation. Mean 100*AUC, accuraccy, and F1-weighted are 99.49, 96.25 % and 96.06 %. To utilize the GLRP, we retrained Graph-CNN on 70 randomly selected samples and applied it on the 8 test samples (4 treated and 4 not treated) to propagate the relevances correspondingly to correctly predicted class. For each out of 8 test samples, we constructed a subnetwork selecting 140 the most relevant genes and deleting singletones, that lead to subnetworks consisting of mainly 130 vertices. Remarkably, the *green* gene module, which was the most strongly correlated one with TNFα upregulation (Rhead et al., 2020) showed significant association (adjusted p-value < 0.05) with the combined set of subnetwork genes, with genes found in the majority of subnetworks and also with 5 of the 8 subnetworks (Supplementary File S1). At the same significance level the *turquoise* gene module described in (Rhead et al., 2020) was strongly associated with 2 of 8 subnetworks and with genes found in all 8 subnetworks. In addition, both the *green* and the *turquoise* modules showed moderate association (adjusted p-value < 0.1) with the majority of gene sets defined on the basis of the test subnetworks. Furthermore, we found strong or moderately significant overlap between upregulated genes and subnetwork gene sets. These results demonstrate partial agreement between gene sets suggested by LRP, another gene network analysis and classical differential expression analysis. Hence, the LRP-based subnetworks gathered biologically meaningful genes and may even complement the other approaches in revealing important properties of the underlying biological systems. Additionally, another two gene sets were compared with WCGNA modules: the intersection of subnetworks genes and genes that occur more than in 4 test samples subnetworks. Notably, the individual subnetworks shared more genes with the *green* and *turquoise* WGCNA modules than those described gene sets, pointing out the ability of GLRP to identify sample-specific genes.

### 3.3 GLRP to deliver patient-specific subnetworks

We applied the developed layer-wise relevance propagation on the Graph-CNN trained on gene expression data from section 2.1. The gene expression data was structured by a protein-protein interaction network. The prediction task was to classify patients into 2 groups, metastatic and non-metastatic. For that, we had two output neurons of the neural network showing the probability of these two classes. The architecture of the Graph-CNN is the same as in our previous study (Chereda et al., 2019) and gene expression data was initially standardized for the training. For the non-image data to standardize the input features is the usual practice. However, in case of standardization, the input features are treated independently. For an image, since the neighboring pixels are highly correlated, if the pixel values are standardized across the dataset this can distort its pattern quite significantly and lead to misinterpretation. Analogically, standardization of microarray data changes expression patterns of genes located in the same neighborhood of a molecular network. This might affect the explainability of the Graph-CNN that we aim at. Therefore, we now train the Graph-CNN directly on the quantile normalized data avoiding the additional standardization step. Instead, we subtracted the minimal value (5.84847) of the data from each cell of the gene expression matrix to keep the gene expression values non-negative. If initially, GE data was lying in [5.84847, 14.2014] now it is in the interval [0.0, 8.3529]. This transformation allows us to apply the LRP propagation rule (7), and to preserve original gene-expression patterns in local neighborhoods of the PPI network. For the comparison we provide the performance of a ‘glmgraph’ method (Chen et al., 2015) implementing network-constrained sparse regression model using HPRD PPI network and Random Forest without any prior knowledge as baselines. glmgraph was not evaluated on non-standardized data, since it has convergence issues in this case. The results of a 10-fold cross validation are depicted in Table 1. The metrics were averaged over folds and the standard errors of their means were calculated.

**Table 1.** Performance of Graph-CNN on metastatic event prediction, depending on normalization.

| Method | Std | 100*AUC | Accuracy, % | F1-weighted, % |
| --- | --- | --- | --- | --- |
| Graph-CNN | - | 82.57±1.25 | 76.07±1.30 | 75.82±1.33 |
| Random Forest | - | 81.27±1.66 | 74.23±1.73 | 73.47±1.84 |
| Graph-CNN | + | 82.16±1.25 | 76.18±1.36 | 75.86±1.35 |
| Random Forest | + | 81.40±1.76 | 74.74±1.67 | 74.00±1.82 |
| glmgraph | + | 80.88±1.37 | 75.14±1.30 | 74.73±1.39 |

The LRP was applied as a post hoc processing. We trained the Graph-CNN on 90% of data, keeping the same hyperparameters from the 10-fold cross validation procedure, and 10% served as a test set. After that, we selected patients from the test set and propagated the relevance only from the Graph-CNN’s output node corresponding to the correctly predicted class. The most frequently selected features are summarized in Supplementary Table S1. The eukaryotic translation elongation factor EEF1A1 which is overexpressed in the majority of breast cancers and protects tumor cells from proteotoxic stress (Lin et al., 2018) was the sole factor that was selected in all of the 97 test set patients. Other frequently selected features in both non-metastatic as well as metastatic patients included genes such as the Epithelial-to-Mesenchymal-Transition (EMT)-related gene VIM (46/58 non-metastatic, 30/39 metastatic patients), the extracellular matrix protein FN1 (43/58 non-metastatic, 22/39 metastatic patients), the actin cytoskeleton regulator CFL1 (7/58 non-metastatic, 7/39 metastatic patients) as well as the estrogen receptor ESR1 28/58 non-metastatic, 10/39 metastatic patients) that are all known to be linked with breast cancer development and progression (Sharma et al., 2019; Wang, Eddy, and Condeelis, 2007; Lin et al., 2019; Feng et al., 2018) This indicates that our method successfully identified relevant key players with a general role in breast tumorigenesis.

Additionally, we show individualized PPI subnetworks delivered for four correctly predicted breast cancer patients (Table 2) from the microarray data set. Two of them had been assigned with the most common subtype luminal A (LumA), while the other two suffered from the highly aggressive basal-like subtype. In each group one patient with early metastasis was picked and one who did not develop any within at least 5 years of follow-up.

**Table 2.** Patients that the PPI subnetworks are generated for.

| Patient's ID | Subtype | Metastatic event | Time of Metastases, years | Last follow-up, years |
| --- | --- | --- | --- | --- |
| GSM519217 | Basal | 1 | 0.9 | - |
| GSM615233 | LumA | 1 | 0.79 | - |
| GSM615695 | Basal | 0 | - | 5.38 |
| GSM150990 | LumA | 0 | - | 9.93 |

The generated PPI subnetworks are displayed in Figure 2. The gene expression level and the relevance score are visualized as the node’s color and size, respectively. The patient-specific subnetworks were created by selecting the PPI’s vertices with the highest relevance scores. The sequence of pictures in order ABCD is the same as in the table.

**Fig. 2.**
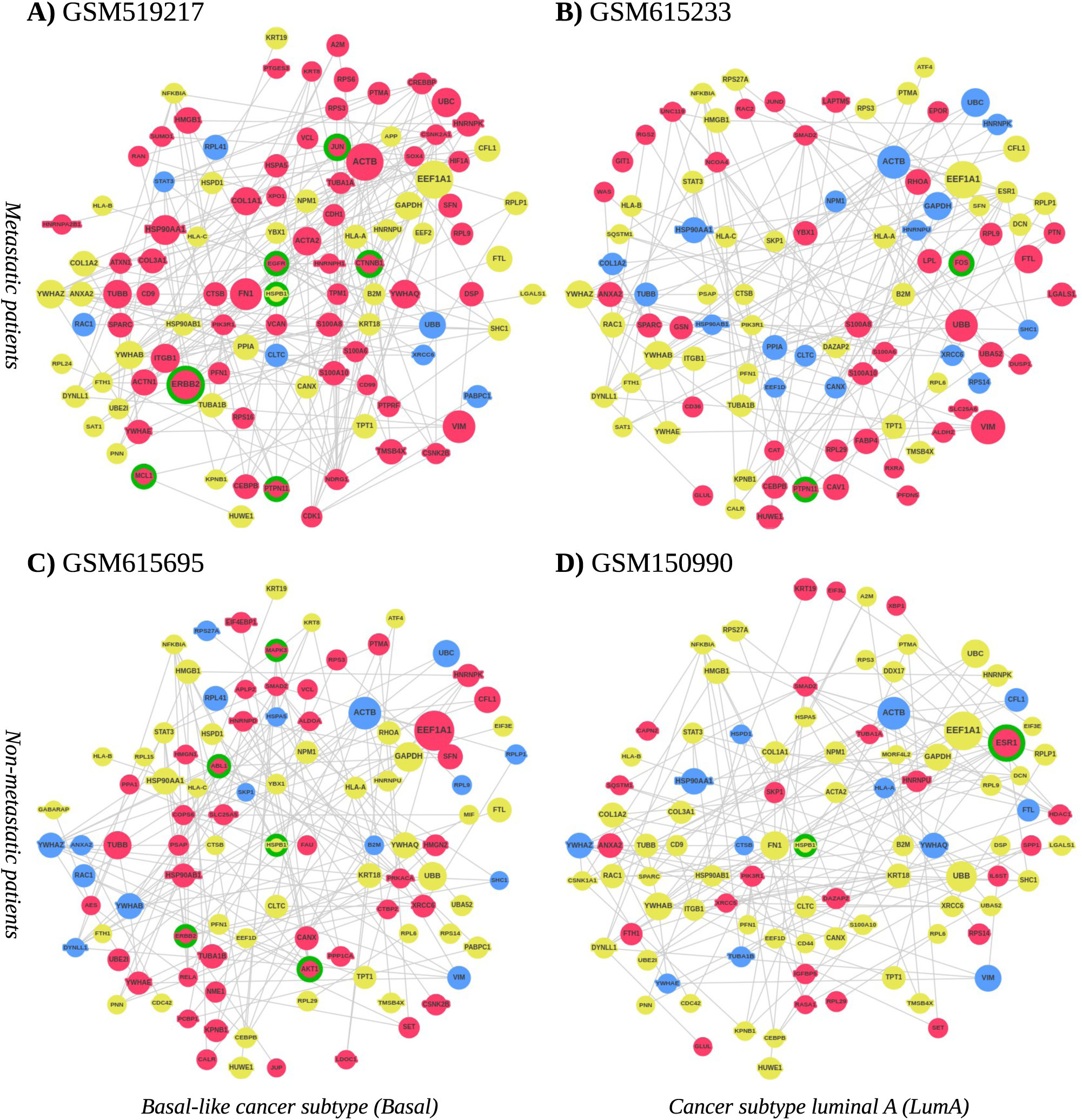
The PPI subnetworks for: 1) Metastatic patients A (GSM519217) and B (GSM615233); 2) Non-metastatic patients C (GSM615695) and D (GSM150990). The coloring of the node is based on gene expression levels by 25% and 75% quantiles (blue=LOW, yellow=NORMAL, red=HIGH). based on the gene expression throughout the whole patient cohort. The size of vertices corresponds to the relevance scores within one subnetwork. All the subnetworks are highly relevant compared to the rest of the PPI network. Green circles highlight targetable genes.

Interestingly, the networks of both LumA patients contained ESR1 which fits well since this subtype is considered as estrogen receptor positive (Perou et al., 2000) In contrast, genes often associated with the basal-like subtype and a poor prognosis such as MCL1, CTNNB1, EGFR or SOX4, were found in the basal-like patient GSM519217 suggesting that the generated networks are capable of extracting breast cancer subtype-specific features. The comparison of the subnetworks of the non-metastatic and the metastatic patients furthermore revealed some patient-specific genes which might give valuable information about specific mechanisms of tumorigenesis and therapeutic vulnerabilities in the respective patient. In general, it seemed that the subnetworks of the non-metastatic patients contained more genes that have been linked to better prognostic outcomes such as JUP, PCBP1 and HMGN2 in GSM615695 (Bailey et al., 2012; Shi et al., 2018; Fan et al., 2018) or RASA1, IL6ST, KRT19 and RPS14 in GSM150990 (Liu et al., 2014; Mathe et al., 2015; Saha et al., 2018; Zhou et al., 2013) while the networks of both metastatic patients harbored genes that are known to be involved in aggressive tumor growth or therapy resistance which might explain the early metastatic spread in these patients. Some examples are CDK1, SFN and XPO1 in GSM519217 (Alexandrou et al., 2019; Neve et al., 2006; Taylor et al., 2019) or CAV1, PTPN11 and FTL in GSM615233 (Qian et al., 2019; Aceto et al., 2012; Chekhun et al., 2013)

However, not only the presence of specific genes might be important, but also their overall expression level. Our analyses identified e.g. the EMT-related gene VIM as one of the most relevant nodes in the subnetworks of both metastatic patients in which the gene was highly expressed (>75% quantile based on the gene expression throughout the whole patient cohort). In contrast, VIM was also present in the subnetworks of the two non-metastatic patients, however, with a lower relevance and a particularly low expression (<25% quantile). VIM is an important marker for EMT and high expression levels correlate with a motile, mesenchymal-like cancer cell state, thus making VIM an essential effector of metastasis (Sharma et al., 2019)

A comparison of subnetwork genes of 79 correctly predicted test set patients to a database of signal transduction pathways confirmed significant enrichment of pathways that have previously been associated with cancer disease mechanisms such as the EGF, ER-alpha, p53 and TGFbeta pathways as well as Caspase and beta-catenin networks. Comparisons were performed for each patient as well as for subtype gene sets formed by combining subnetwork genes of patients associated with a breast cancer subtype. Results for the 238 signaling pathways from the TRANSPATH^®^ database that were significantly enriched with subtype genes are visualized in Figure 3. Differences in enrichment significance may suggest that the importance of some signaling pathways detected this way is subtype-specific, e.g. for YAP ubiquitination or the VE-cadherin network (orange heatmap, Fig. 3, see also Supplementary Table S2 for details). The pattern of enrichment found on the level of cancer subtypes coincided well with the findings for subnetwork genes of individual patients revealing several molecular networks with elevated significance in both subtype and patient gene sets such as the EGF pathway, although the patient-level visualization did not suggest subtype-specific enrichment (green heatmap, Fig. 3). One source of these observations can be that patient subnetworks tend to be associated with certain pathways but cover different pathway components (genes). We therefore compared pathway genes in pairs of patient subnetworks for the 33 largest pathways. In 18 pathways the median pair of patient subnetworks differed in 33% or more of the genes matched within a pathway (see also Supplementary Table S3 for details). These results demonstrate that the subnetworks obtained by Graph-CNN were enriched with common signaling pathways relevant for the respective disease and can assign patient-specific priorities to pathway components.

**Fig. 3.**
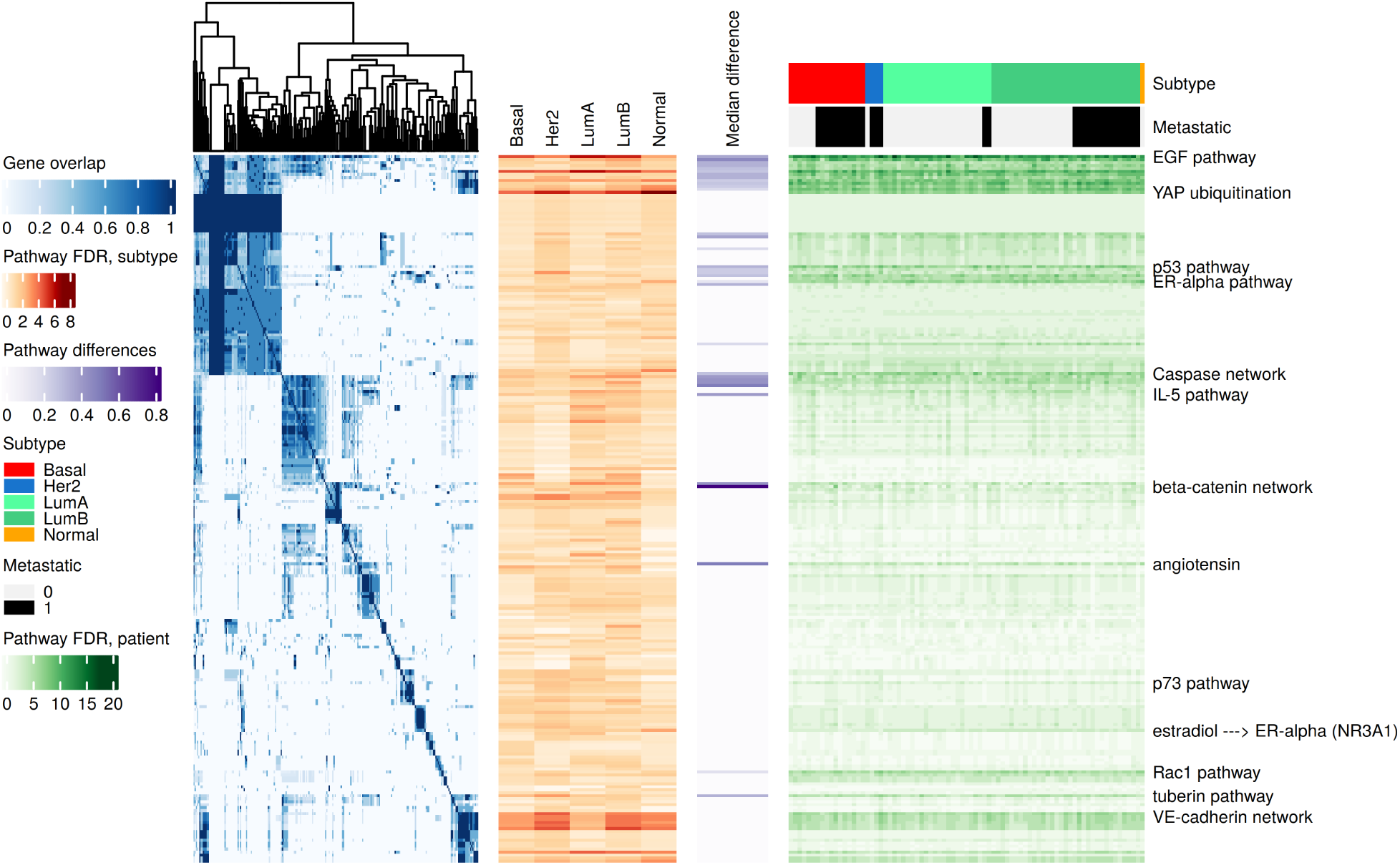
Signal transduction pathway analysis of subnetwork genes reported for 79 patients in 5 subtypes. (From left to right) Blue heatmap: 238 signaling pathways clustered according to proportion of shared subnetwork genes; Orange heatmap: Enrichment significance of pathways in subnetwork genes combined from patients of given subtype. Darker orange indicates higher significance; Purple heatmap: Median difference in matched pathway genes observed in pairwise comparisons of subnetwork gene sets from patients mapped to 33 pathways. Darker purple indicates higher tendency of pairs of subnetwork gene sets to coincide with different pathway genes; Green heatmap: Enrichment significance of pathways in subnetwork genes of 79 patients. Darker green indicates higher significance. Corresponding subtypes and metastatic status are shown by the annotation above the heatmap. A detailed version of this figure capturing pathway and sample names is provided in Supplementary Figure S2.

Finally, we tested whether the subnetworks can also be used for finding potentially targetable genetic vulnerabilities that could open new options for personalized treatment decisions. We applied the “MTB report” methodology described in (Perera-Bel et al., 2018) to identify actionable genes present in the subnetworks. For that, we extended the algorithm to match high expression with gain of function alterations, and low expression with loss of function alterations. The results are summarized in Table 3.

**Table 3.**
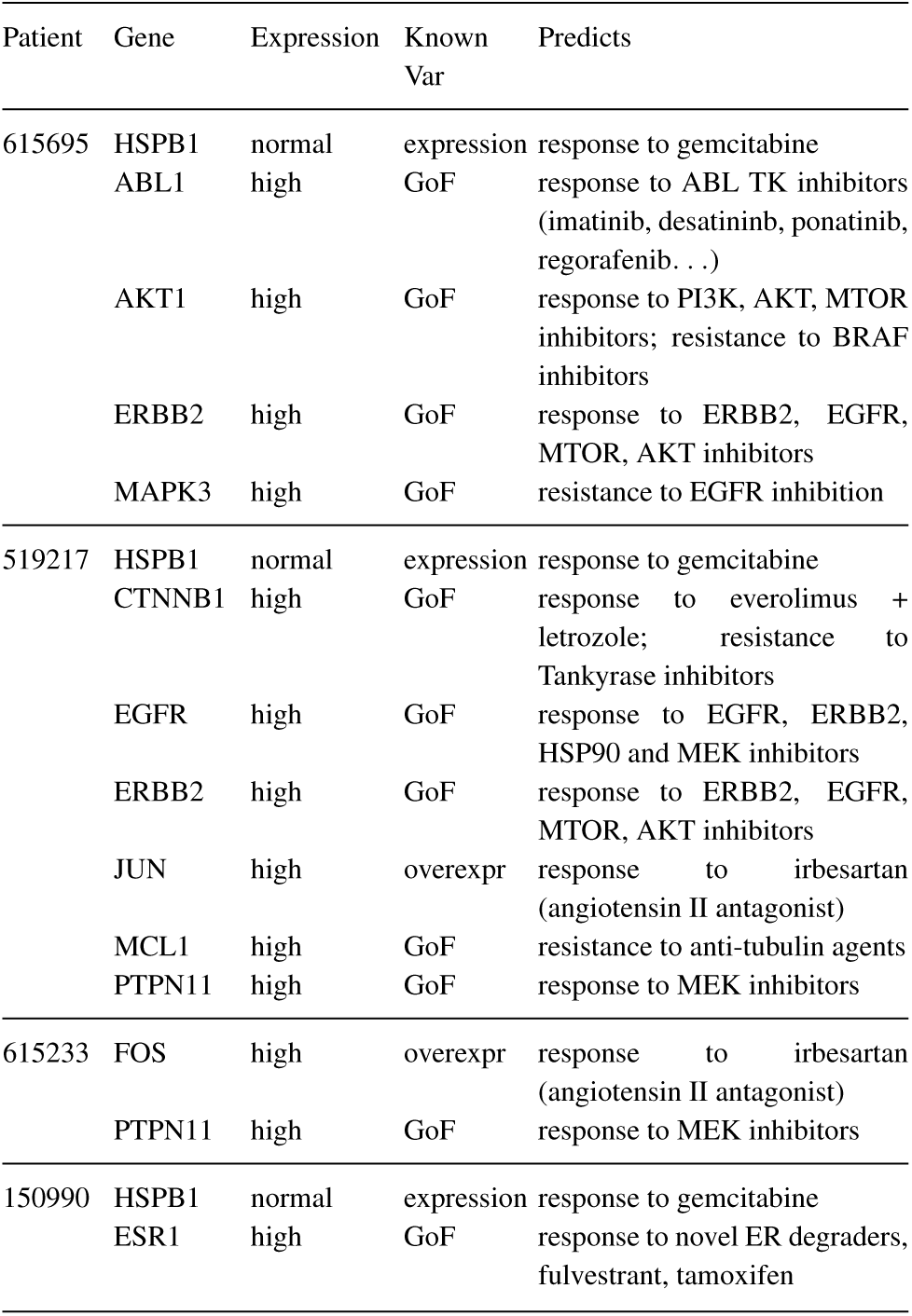
Actionable genes identified by the MTB Report workflow. Genes from the PPI subnetworks were matched to known genomic alterations (Known Var) that predict either response or resistance to drugs (Predicts). High and low gene expression were matched to gain of function (GoF) and loss of function (LoF) genomic variants, respectively.

Although information about the presence of actionable genetic variants is missing from our patient microarray data, the information generated by the PPI subnetworks could be used to define specific panels for subsequent sequencing. Indeed, the MTB reports highlighted specific genes that could be targeted therapeutically in each of the four patients: In the non-metastatic LumA patient GSM150990 ESR1 was proposed as therapeutic target which is in line with current treatment regimens that use hormone therapy as the main first line treatment of choice for this patient subgroup. In contrast, in the metastatic LumA patient GSM615233 FOS and PTPN11 were identified as novel actionable alterations. In the often rapidly-relapsing basal-like patients HSPB1 and ERBB2 were identified as common targets as well as MAPK3, AKT1 and ABL1 for the non-metastatic patient GSM615695 or EGFR, MCL1, CTNNB1, PTPN11 and JUN for the metastatic patient GSM519217, thereby suggesting novel possibilities for combinatory or alternative treatments. Taken together, GLRP provides subnetworks centered around known oncogenic drivers that seem reasonable in the context of cancer biology and can help to identify patient-specific cancer dependencies and therapeutic vulnerabilities in the context of precision oncology.

## 4 Discussion

In our work we focused on the interpretability of a deep learning method utilizing molecular networks as prior knowledge. We implemented LRP for Graph-CNN and provided the sanity check of the developed approach on the MNIST dataset. Essentially, the main aim of the paper was to explain the prediction of metastasis for breast cancer patients by providing an individual molecular subnetwork specific for each patient. The patient-specific subnetworks provided interpretability of the deep learning method and demonstrated clinically relevant results on the breast cancer dataset.

Supposedly, the performance of Graph-CNN can be improved. The batch normalization technique (Ioffe and Szegedy, 2015) that is used to accelerate the training of deep neural networks is not seen to be available for the Graph-CNN, so this can be the way to enhance its performance. The LRP rule for batch normalization layers is yet another procedure to be adapted for Graph-CNN.

Another possibility to identify genes (and construct subnetworks out of them) influencing classifier decisions is to apply model-agnostic SHAP and LIME explainability methods. LIME method provides explanations of a data-point based on feature perturbations. The method samples perturbations from a Gaussian distribution, ignoring correlations between features. It leads to the instability of explanations that is not favourable for personalised medicine. SHAP provides Shapley values for each feature of a data-point as well but does not have such an issue, so we attempted to derive patients-specific subnetworks applying TreeExplainer and KernelExplainer from SHAP python module on Random Forest and Graph-CNN respectively. The subnetworks were build on the basis of HPRD PPI utilizing positive Shapley values, which were pushing prediction to a higher probability of corresponding class (metastatic or non-metastatic). The subnetworks obtained are mostly consisting from single vertices. In contrast, the subnetworks from GLRP and Graph-CNN are mostly connected. The SHAP’s DeepExplainer approach suitable for convenient deep learning models is not applicable for Graph-CNN. The model-agnostic KernelExplainer computes SHAP values out of a debiased lasso regression. Reevaluating the model happens several thousands numbers of times specified by a user as well as a small background dataset needed for integrating out features. Hence, the KernelExplainer is not scalable and application of it on Graph-CNN resulted in not connected subnetworks as well.

Furthermore, the sensitivity of Graph-CNN to the changes of prior knowledge is still to be investigated. Authors in (Defferrard, Bresson, and Vandergheynst, 2016) showed that for the MNIST images a random graph connecting pixels significantly decreases the performance destroying local connectivity. In our case, the permutation of the vertices of the PPI network does not influence the classifier performance on standardized gene expression data. Yet, PPI network is a small world network and its degree distribution fits to the power law with the exponent *α* = 2.70. It implies great connectivity between proteins and means that any two nodes are separated by less than six hops. The filters of convolutional layers cover a 7-hop neighborhood of each vertex, so we assume it still might be enough to capture the gene expression patterns. In our future work we will investigate how the properties of the prior knowledge influence the performance and explainability of Graph-CNN.

The subnetworks generated by GLRP contained common potential oncogenic drivers which indicates that they can extract the essential cancer pathways. Indeed, our analyses identified genes associated with hormone receptor positive breast cancer (e.g. ESR1, IL6ST, CD36, GLUL, RASA1) in the networks from the patients with estrogen receptor positive, LumA breast cancer and genes associated with the basal-like subtype (e.g. EGFR, SOX4, AKT1 as well as high levels of HNRNPK) in the basal-like patients, underlining the biological relevance of the networks. Next to subtype-specific genes, the networks contained several oncogenes that were found in all four patients and could thus represent common drivers of breast cancer initiation and progression. One example is the actin-binding protein cofilin (CFL1) that regulates cancer cell motility and invasiveness (Wang, Eddy, and Condeelis, 2007) Another interesting candidate is STAT3 which is activated in more than 40% of breast cancers and can cause deregulated cell proliferation and Epithelial-to-Mesenchymal Transition (EMT) (Banerjee and Resat, 2015) Our graphs not only displayed patient-specific PPI subnetworks, but also concisely visualized the relevance of each node and its expression levels. This information is potentially relevant to judge the biological significance of the gene in a patient-specific context.

Next to the common genes found in all four networks, each network was characterized by several special, cancer-associated genes which are of high interest because they might represent patient-specific central signaling nodes and therapeutic vulnerabilities. Some examples are PTPN11 that is known to activate a transcriptional program associated with cancer stem cells or the EMT-related genes SOX4 or VIM that might be responsible for the high invasive capacity of the tumors and their early metastasis formation (Bentires-Alj et al., 2004; Aceto et al., 2012; Sharma et al., 2019; Zhang et al., 2012) Interestingly, the network of the metastatic patient GSM615233 harbored the genes FABP4 and LPL which both have been shown to interact with CD36, another highly expressed node in the network, to support cell proliferation and counteract apoptosis (Guaita-Esteruelas et al., 2016; Liang et al., 2018; Kuemmerle et al., 2011) In contrast, in the non-metastatic patient GSM150990 especially the interleukin receptor IL6ST and the Ras GTPase-activating protein 1 (RASA1) seem to be interesting because for both high expression levels have been linked with a favorable prognosis (Liu et al., 2014; Mathe et al., 2015) In the other non-metastatic patient GSM615695 high levels of HMGN2 and PCBP1 were identified which both have been shown to be able to inhibit cell proliferation (Shi et al., 2018; Fan et al., 2018) Although the experimental validation for the networks is still missing, it is tempting to speculate that these genes might contribute to the benign phenotype of the tumor in these patients.

All patient-specific subnetworks contained relevant drug targets that have been largely studied in breast cancer (e.g. ERBB2, ESR1, EGFR, AKT1). Yet, resistance mechanisms in breast cancer targeted therapies represent a big challenge; many of the identified therapeutic approaches have failed (Nakai, Hung, and Yamaguchi, 2016) due to the highly interconnected nature of signaling pathways and potential circumvents. A promising way forward could involve the molecular characterization of the tumor with transcriptomics and a parallel culture of patient-derived organoids. PPI networks could elucidate the right combination strategy by identifying central signaling nodes. Different therapeutic strategies could be tested on organoids and confirm the best strategy that synergistically blocks cancer cell escape routes and minimizes the emergence of survival mechanisms. Only the identification of relevant mechanisms of action for cell survival as well as of the factors involved in resistance for each patient, together with a more precise and personalized characterization of each cancer phenotype, may provide useful improvements in current therapeutic approaches.

## 5 Conclusion

We present a novel Graph-CNN based feature selection method that benefits from prior knowledge and provides patient-specific subnetworks. We adapted the existing Layer-wise Relevance Propagation technique to the Graph-CNN, demonstrated it on MNIST data, and showed its applicability on a large breast cancer dataset. Our new approach generated individual patient-specific molecular subnetworks that influenced the model’s decision in the given context of a classification problem. The subnetworks selected by the developed method utilizing general prior knowledge are relevant for prediction of metastasis in breast cancer. They contain common as well as subtype-specific cancer genes that match the clinical subtype of the patients, together with patient-specific genes that could potentially be linked to aggressive/benign phenotypes. In the context of a breast cancer dataset GLRP provides patient-specific explanations for the Graph-CNN that largely agree with clinical knowledge, include oncogenic drivers of tumor progression and can help to identify therapeutic vulnerabilities. We therefore conclude that our method GLRP in combination with Graph-CNN is a new, useful and interpretable machine learning approach for high-dimensional genomic data-sets. Generated classifiers rely on prior knowledge of molecular networks and can be interpreted by patient-specific subnetworks driving the individual classification result. These sub-networks can be visualized and interpreted in a biomedical context on the individual patient level. This approach could thus be useful for precision medicine approaches such as for example the molecular tumorboard.

## Supporting information

**Supplementary File S1. Subnetwork genes obtained for 8 test samples and analysis of their association with gene modules reported by (Rhead et al., 2020) as well as differentially expressed (DE) genes.** Worksheet *Subnetwork genes 8 samples* provides identifiers and gene symbols of 167 subnetwork genes, in how many and in which samples they were selected. Worksheet *Gene module enrichment* presents results of Fisher test calculations comparing subnetwork gene sets to gene modules and DE gene sets. Each row contains data for a DE gene set or a gene module consisting of the total group size and column tripletts with p-value, adjusted p-value as well as the number of hits, respectively, observed in comparisons to the union of genes from 8 subnetworks, the set of genes occurring in the majority, the set of genes found in all of the subnetworks and each of the 8 samples. Highlighted are rows corresponding to *green* and *turquoise* gene modules, which were most often significantly associated with subnetwork gene sets (grey), adjusted p-values below 0.05 (red) and between 0.05 and 0.1 (yellow).

**Supplementary Figure S1. Visualization of 2 out of 32 learned filters of the 1st convolutional layer of graph CNN classifying MNIST digits on the 8-nearest-neighbours graph.**

**Supplementary Figure S2. Signal transduction pathway analysis of subnetwork genes reported for 79 correctly classified test set patients in 5 subtypes.**

**Supplementary Table S1. Frequency of gene selection in top 10 of highly relevant genes among metastatic and non-metastatic patients.**

**Supplementary Table S2. 238 signal transduction pathways from the TRANSPATH**^**®**^ **database that were significantly enriched (FDR < 0.05) in subnetwork genes associated with 5 cancer subtypes.**

## Acknowledgements

This work was funded by the German Ministry of Education and Research (BMBF) e:Med project *MyPathSem* (031L0024) and the project *MTB-Report* by the big data initiative of the Volkswagenstiftung. K.M. was supported by German Research Foundation (DFG) project 424252458. H.C. is a member of the International Max Planck Research School for Genome Science, part of the Göttingen Graduate Center for Neurosciences, Biophysics, and Molecular Biosciences. T.B. is a member of the Göttingen Campus Institute Data Science. We would like to acknowledge Michaela Bayerlová for fruitful discussions.

