## Supplementary_Figures_and_Tables for "Explaining decisions of Graph Convolutional Neural Networks: patient-specific molecular subnetworks responsible for metastasis prediction in breast cancer"

### Supplementary Figure S1

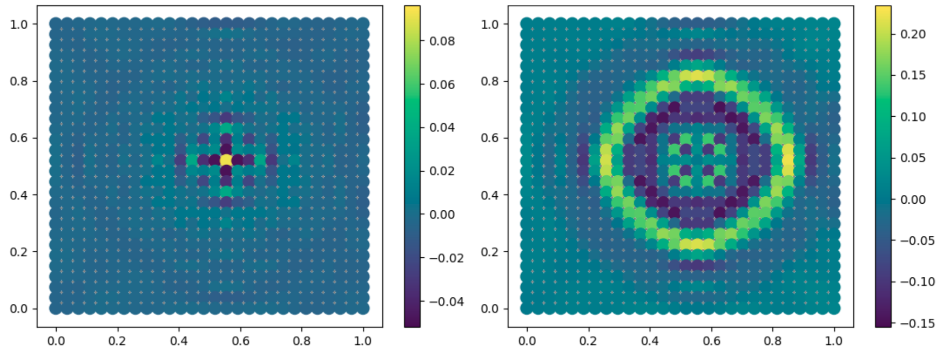

**Fig. 1**. Visualization of 2 out of 32 learned filters of the 1st convolutional layer of graph CNN classifying MNIST digits on the 8-nearest-neighbours graph. The convolutional filters have 25 parameters each. This means that the neighborhood covered by convolution can be expanded up to 24 hops from the vertice where the filter is centered. Due to rotational invariance, filters tend to learn patterns that concern rounded curves on the 8-nearest-neighbours graph. PyGSP[1] package was used for visualization.

### Supplementary Table S1

**Table 1.** Frequency of gene selection in top 10 of highly relevant genes among metastatic and non-metastatic patients.

| Metastatic (39 patients) | | Non-metastatic (58 patients) | |
|---|---|---|---|
| **Gene** | **Frequency** | **Gene** | **Frequency** |
| EEF1A1 | 39 | EEF1A1 | 58 |
| UBC | 39 | ACTB | 57 |
| ACTB | 39 | UBB | 57 |
| UBB | 38 | UBC | 56 |
| GAPDH | 33 | HSP90AA1 | 46 |
| HSP90AA1 | 31 | VIM | 46 |
| VIM | 30 | GAPDH | 44 |
| YWHAQ | 29 | FN1 | 43 |
| YWHAZ | 29 | YWHAB | 42 |
| YWHAB | 27 | YWHAQ | 30 |
| FN1 | 22 | ESR1 | 28 |
| ESR1 | 10 | YWHAZ | 25 |
| TUBB | 8 | RPL41 | 18 |
| FTL | 7 | TUBB | 10 |
| CFL1 | 7 | COL1A2 | 9 |
| RPL41 | 6 | COL1A1 | 9 |
| PPIA | 5 | FTL | 8 |
| ERBB2 | 5 | CFL1 | 7 |
| COL1A1 | 4 | CLTC | 7 |
| KRT18 | 4 | KRT18 | 7 |

[1] Defferrard, M., Martin, L., Pena, R., and Perraudin, N. PyGSP: Graph Signal Processing in Python. DOI: 10.5281/zenodo.1003157

### Supplementary Figure S2

**Figure 2.** Signal transduction pathway analysis of subnetwork genes reported for 79 correctly classified test set patients in 5 subtypes. This image resembles the corresponding figure of the main text while diplaying also pathway and sample names. The visualization was created using the ComplexHeatmap R package[2]. (From left to right) Blue heatmap: 238 signaling pathways clustered according to proportion of shared subnetwork genes; Orange heatmap: Enrichment significance of pathways in subnetwork genes combined from patients of given subtype. Darker orange indicates higher significance; Purple heatmap: Median difference in matched pathway genes observed in pairwise comparisons of subnetwork gene sets from patients mapped to 33 pathways. Darker purple indicates higher tendency of pairs of subnetwork gene sets to coincide with different pathway genes; Green heatmap: Enrichment significance of pathways in subnetwork genes of 79 patients. Darker green indicates higher significance. Corresponding subtypes and metastatic status are shown by the annotation above the heatmap.

[2] Gu, Z., Eils, R., Schlesner, M. (2016). Complex heatmaps reveal patterns and correlations in multidimensional genomic data, *Bioinformatics*, 32(18), 2847–2849, https://doi.org/10.1093/bioinformatics/btw313.

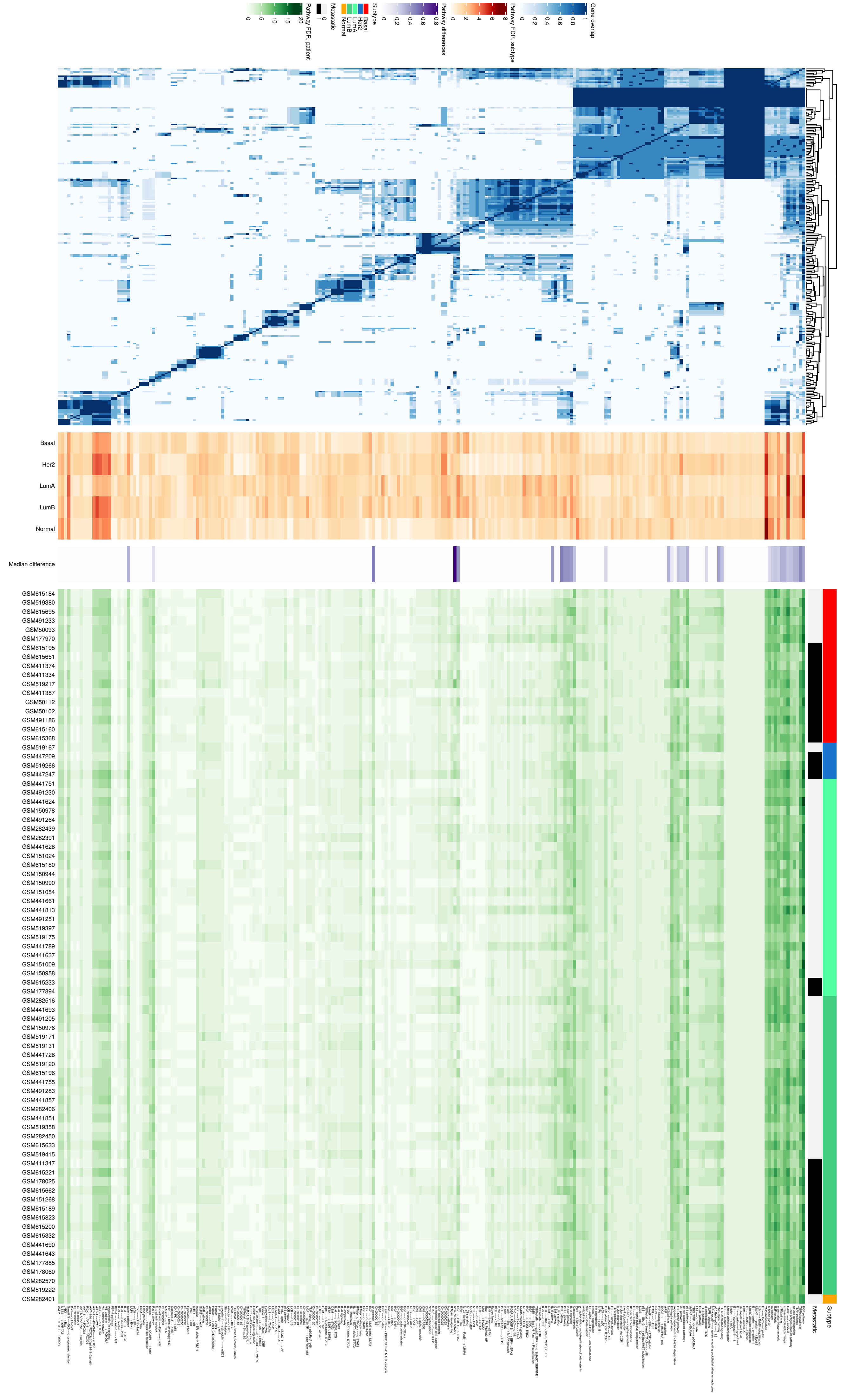

Gene overlap
0
0.2
0.4
0.6
0.8
1
Pathway FDR, subtype
0
2
4
6
8
Pathway differences
0
0.2
0.4
0.6
0.8
Subtype
Basal
Her2
LumA
LumB
Normal
Metastatic
0
1
Pathway FDR, patient
0
5
10
15
20
Basal
Her2
LumA
LumB
Normal
Median difference
GSM615184
GSM519380
GSM615695
GSM491233
GSM50093
GSM177970
GSM615195
GSM615651
GSM411374
GSM411334
GSM519217
GSM411387
GSM50112
GSM50102
GSM491186
GSM615160
GSM615368
GSM519167
GSM447209
GSM519266
GSM447247
GSM441751
GSM491230
GSM441624
GSM150978
GSM491264
GSM282439
GSM282391
GSM441626
GSM151024
GSM615180
GSM150944
GSM150990
GSM151054
GSM441661
GSM441813
GSM491251
GSM519397
GSM519175
GSM441789
GSM441637
GSM151009
GSM150958
GSM615233
GSM177894
GSM282516
GSM441693
GSM491205
GSM150976
GSM519171
GSM519131
GSM441726
GSM519120
GSM615196
GSM441755
GSM491283
GSM441857
GSM282406
GSM441851
GSM519358
GSM282450
GSM615633
GSM519415
GSM411347
GSM615221
GSM178025
GSM615662
GSM151268
GSM615189
GSM615823
GSM615200
GSM615332
GSM441690
GSM441643
GSM177885
GSM178060
GSM282570
GSM519222
GSM282401
Metastatic
Subtype

### Supplementary Table S2

**Table 2.** 238 signal transduction pathways from the TRANSPATH® database that were significantly enriched (FDR < 0.05) in subnetwork genes associated with 5 cancer subtypes. The table shows pathway accessions and names as well as the -$\log_{10}$(adjusted p-value) of Fisher's exact test which was used to determine enrichment of pathway genes in subtype gene sets. Adjustment of p-values was carried out using the Benjamini-Hochberg method[3]. Transformation of values was performed to support display in heatmaps. The sum was calculated to sort pathways by their overall importance.

| Pathway | Name | Basal | Her2 | LumA | LumB | Normal | Sum |
|---|---|---|---|---|---|---|---|
| CH000004693 | YAP ubiquitination | 4.174 | 5.528 | 4.38 | 5.153 | 6.821 | 26.056 |
| CH000004191 | ErbB3 ---> survival | 4.692 | 4.437 | 5.925 | 5.171 | 3.916 | 24.141 |
| CH000000722 | EGF pathway | 4.692 | 3.649 | 5.851 | 5.453 | 2.944 | 22.588 |
| CH000004208 | AKT-1 ---/YAP | 3.513 | 4.617 | 2.712 | 4.827 | 3.916 | 19.584 |
| CH000004646 | Smad2/3 ---TAZ---> cytoplasmic retention | 3.091 | 3.206 | 4.38 | 4.005 | 3.589 | 18.271 |
| CH000004615 | ANG-1 ---/ FOXO1A | 3.091 | 4.437 | 2.472 | 3.943 | 3.098 | 17.041 |
| CH000004598 | VE-cadherin network | 3.093 | 3.726 | 2.647 | 3.943 | 3.57 | 16.978 |
| CH000004597 | VE-cadherin ---> FOXO1A | 3.093 | 3.726 | 2.647 | 3.943 | 3.57 | 16.978 |
| CH000004613 | Angiopoietin/Tie signaling | 3.006 | 4.437 | 2.472 | 3.64 | 3.357 | 16.912 |
| CH000004539 | AKT-1 ---PRAS40---> mTOR | 3.006 | 4.086 | 2.285 | 3.902 | 3.57 | 16.849 |
| CH000004600 | mammalian Hippo network | 3.025 | 2.893 | 3.142 | 2.854 | 3.647 | 15.561 |
| CH000000692 | S phase (Cdk2) | 2.302 | 2.695 | 2.053 | 2.661 | 3.57 | 13.281 |
| CH000000321 | CH000000321 | 2.725 | 3.532 | 2.718 | 2.668 | 1.42 | 13.063 |
| CH000000322 | CH000000322 | 2.725 | 3.532 | 2.718 | 2.668 | 1.42 | 13.063 |
| CH000004475 | IL-3 signaling | 2.409 | 1.767 | 3.017 | 3.236 | 2.292 | 12.721 |
| CH000004599 | MST2 ---> YAP, TAZ | 1.89 | 2.695 | 2.033 | 2.632 | 3.355 | 12.604 |
| CH000004572 | EGF ---Raf-1---> ERK2 | 2.789 | 1.639 | 3.496 | 3.236 | 0.938 | 12.1 |
| CH000004526 | autophagy | 1.77 | 2.414 | 1.584 | 2.603 | 3.35 | 11.72 |
| CH000000580 | Fam ---> beta-catenin | 2.074 | 1.488 | 2.285 | 2.147 | 3.57 | 11.563 |
| CH000000711 | TGFbeta pathway | 1.995 | 1.099 | 2.647 | 3.065 | 1.962 | 10.767 |
| CH000000743 | tuberin pathway | 2.406 | 3.148 | 2.053 | 1.838 | 1.279 | 10.724 |
| CH000000756 | OSM pathway | 1.259 | 1.639 | 2.891 | 2.665 | 1.872 | 10.326 |
| CH000004192 | NRG ---> ERK | 1.944 | 1.347 | 3.311 | 2.537 | 1.085 | 10.224 |
| CH000004538 | AMPK ---14-3-3---/ mTOR | 1.526 | 2.2 | 1.621 | 2.026 | 2.804 | 10.177 |
| CH000000626 | EGF ---> STAT1alpha, STAT3 | 2.725 | 2.2 | 1.791 | 2.668 | 0.765 | 10.149 |
| CH000003550 | sumoylation pathway | 1.794 | 2.434 | 1.925 | 2.661 | 1.179 | 9.993 |
| CH000000232 | CH000000232 | 1.89 | 2.414 | 2.018 | 1.962 | 1.487 | 9.77 |
| CH000004506 | IL-5 pathway | 2.14 | 1.639 | 2.366 | 2.147 | 1.392 | 9.685 |
| CH000004243 | estradiol ---> ER-alpha (NR3A1) | 1.187 | 1.675 | 1.925 | 1.832 | 2.95 | 9.569 |
| CH000000879 | Caspase network | 1.989 | 1.828 | 1.596 | 2.661 | 1.487 | 9.561 |
| CH000004625 | MIC2 signaling | 3.006 | 0.866 | 1.621 | 2.884 | 1.002 | 9.379 |
| CH000000736 | PDGF pathway | 1.89 | 1.512 | 1.721 | 3.003 | 1.179 | 9.304 |
| CH000000956 | E1 ---/ alpha-tubulin | 1.522 | 2.127 | 1.621 | 1.49 | 2.481 | 9.24 |
| CH000004667 | MEK ---> EZR | 1.526 | 0.62 | 2.472 | 2.374 | 2.217 | 9.21 |
| CH000000987 | hypoxia pathways | 1.372 | 1.042 | 2.033 | 3.08 | 1.664 | 9.19 |
| CH000004634 | TGFbetaR-I ---> ERK | 1.693 | 0.617 | 2.975 | 2.171 | 1.539 | 8.995 |
| CH000000869 | p73 pathway | 1.921 | 2.185 | 1.596 | 1.962 | 1.279 | 8.942 |
| CH000004649 | AT1A ---> STAT1, STAT3 | 1.357 | 1.871 | 2.285 | 2.147 | 1.279 | 8.938 |
| CH000004583 | homophilic ligation of E-cadherin | 1.906 | 1.767 | 2.053 | 1.965 | 1.211 | 8.902 |

[3] Benjamini, Y., & Hochberg, Y. (1995). Controlling the False Discovery Rate: A Practical and Powerful Approach to Multiple Testing. *Journal of the Royal Statistical Society. Series B (Methodological), 57*(1), 289-300.

| | | | | | | | |
|---|---|---|---|---|---|---|---|
| CH000001012 | ARIP1 ---> atrophin1 | 1.921 | 1.635 | 2.093 | 1.962 | 1.279 | 8.889 |
| CH000000755 | TLR4 pathway | 1.666 | 1.899 | 1.508 | 2.276 | 1.487 | 8.836 |
| CH000000123 | CH000000123 | 1.693 | 2.2 | 1.791 | 1.729 | 1.42 | 8.834 |
| CH000000124 | CH000000124 | 1.693 | 2.2 | 1.791 | 1.729 | 1.42 | 8.834 |
| CH000001016 | AR pathway | 1.187 | 3.251 | 1.654 | 1.43 | 1.111 | 8.633 |
| CH000004590 | tubulin ---restin, IQGAP1---> actin | 1.315 | 2.165 | 1.508 | 1.729 | 1.909 | 8.626 |
| CH000000999 | EGF ---> c-Fos | 1.522 | 0.866 | 3.005 | 2.147 | 1.002 | 8.542 |
| CH000000721 | SMRT ---/ AhR | 1.794 | 2.434 | 1.925 | 1.2 | 1.179 | 8.532 |
| CH000000851 | JNK pathway | 1.121 | 2.695 | 1.288 | 1.867 | 1.487 | 8.458 |
| CH000003912 | neurotrophic signaling | 1.581 | 0.962 | 1.791 | 2.388 | 1.586 | 8.307 |
| CH000000030 | EGF ---> ERK1, ERK2 | 1.574 | 0.9 | 2.399 | 2.264 | 1.058 | 8.195 |
| CH000004510 | IL-5 ---> ERK | 1.234 | 1.447 | 1.791 | 2.171 | 1.539 | 8.183 |
| CH000004533 | STAT1 ---IRF-1---> IRF2 | 1.693 | 2.2 | 1.791 | 1.729 | 0.765 | 8.179 |
| CH000000724 | EGF ---> ERK | 1.526 | 1.114 | 2.216 | 2.026 | 1.279 | 8.161 |
| CH000000719 | AhR pathway | 1.992 | 2.2 | 1.621 | 1.49 | 0.831 | 8.134 |
| CH000000926 | PRL pathway | 1.89 | 1.365 | 1.68 | 2.171 | 0.914 | 8.019 |
| CH000003878 | SOCS-1 ---> p50:RelA-p65 | 1.526 | 1.995 | 1.621 | 1.514 | 1.328 | 7.985 |
| CH000004521 | TLR2-mediated signaling | 1.023 | 2.342 | 1.171 | 2.232 | 1.179 | 7.947 |
| CH000000715 | HIF-1alpha pathway | 1.526 | 1.709 | 1.459 | 2.026 | 1.179 | 7.899 |
| CH000000747 | E2F network | 1.77 | 1.675 | 1.038 | 1.884 | 1.529 | 7.896 |
| CH000004623 | MIC2-isoform2 ---FosB---> MMP9 | 2.693 | 0.516 | 1.291 | 2.661 | 0.626 | 7.787 |
| CH000000718 | HIF-1alpha ---/ AhR | 1.794 | 1.675 | 1.291 | 1.832 | 1.179 | 7.771 |
| CH000000583 | Siah ---> beta-catenin | 1.12 | 1.767 | 1.243 | 1.49 | 2.111 | 7.731 |
| CH000004471 | IFNgamma signaling | 1.061 | 2.2 | 1.584 | 1.399 | 1.42 | 7.664 |
| CH000000839 | c-Jun degradation via COP1 | 1.239 | 1.767 | 1.375 | 1.292 | 1.909 | 7.58 |
| CH000004341 | ER-alpha pathway | 0.807 | 1.639 | 1.14 | 1.162 | 2.804 | 7.553 |
| CH000000750 | insulin pathway | 1.526 | 1.393 | 2.033 | 1.399 | 1.179 | 7.53 |
| CH000000770 | beta-catenin network | 1.855 | 1.531 | 1.776 | 1.729 | 0.615 | 7.505 |
| CH000000874 | ERK1 ---/ Tau | 1.234 | 0.767 | 2.366 | 2.298 | 0.837 | 7.503 |
| CH000003547 | PIAS2-alpha ---SUMO-1---/ AR | 0.991 | 2.2 | 1.791 | 1.729 | 0.765 | 7.476 |
| CH000004149 | Nrf2 pathway | 1.006 | 1.365 | 1.508 | 2.068 | 1.51 | 7.456 |
| CH000000758 | stress-associated pathways | 1.186 | 1.773 | 1.338 | 1.492 | 1.632 | 7.421 |
| CH000000045 | Hsp90 ---> AhR | 1.357 | 1.871 | 1.508 | 1.399 | 1.279 | 7.414 |
| CH000000909 | Ubc9 ---> AP-2 | 1.52 | 1.871 | 1.596 | 1.514 | 0.904 | 7.404 |
| CH000004666 | NF-kappaB ---> genes encoding endothelial adhesion molecules | 0.991 | 2.165 | 1.086 | 1.729 | 1.42 | 7.391 |
| CH000000997 | Ubc9 ---/ p73alpha | 1.526 | 1.995 | 1.621 | 1.514 | 0.73 | 7.387 |
| CH000004484 | KSR scaffold complex | 1.466 | 1.635 | 1.596 | 1.407 | 1.279 | 7.383 |
| CH000000603 | Epo ---> STAT1alpha, STAT3 | 1.693 | 1.365 | 1.791 | 1.729 | 0.765 | 7.343 |
| CH000003552 | SUMO-1 ---> Daxx ---/ CBP | 1.89 | 1.488 | 1.171 | 1.962 | 0.792 | 7.303 |
| CH000004455 | caveolin-1 ---calmodulin1---> eNOS | 1.693 | 1.635 | 1.186 | 1.122 | 1.644 | 7.28 |
| CH000004722 | cxcr4 ubiquitination | 1.693 | 1.365 | 1.065 | 1.729 | 1.42 | 7.272 |
| CH000000952 | UCH-L1 ---> alpha-synuclein | 1.079 | 1.488 | 1.171 | 1.962 | 1.487 | 7.186 |
| CH000000624 | EGF ---/ AKT | 1.239 | 1.767 | 1.375 | 1.965 | 0.655 | 7 |
| CH000004624 | MIC2-isoform2 ---JNK, JunD---> MMP9 | 2.074 | 1.157 | 0.851 | 2.147 | 0.695 | 6.924 |
| CH000004477 | IL-3 ---> 14-3-3 | 1.22 | 1.257 | 1.813 | 1.217 | 1.41 | 6.918 |
| CH000004247 | ER-alpha (NR3A1) ubiquitination | 0.814 | 1.157 | 1.508 | 1.399 | 2.039 | 6.917 |
| CH000000573 | phosphorylation and ubiquitination of beta-catenin | 1.061 | 1.257 | 1.171 | 1.399 | 2.014 | 6.903 |
| CH000000576 | AKT, MAPKAPK2 ---/ tuberin | 1.693 | 1.773 | 1.334 | 1.217 | 0.868 | 6.886 |

| | | | | | | | |
|---|---|---|---|---|---|---|---|
| CH000000663 | Hsp90 ---> AhR (CH000000663) | 1.239 | 1.767 | 1.375 | 1.292 | 1.211 | 6.883 |
| CH000000725 | c-Cbl ---/ ErbB1 | 1.239 | 1.767 | 1.375 | 1.292 | 1.211 | 6.883 |
| CH000000761 | IL-10 pathway | 0.814 | 1.871 | 1.508 | 1.399 | 1.279 | 6.87 |
| CH000000584 | IL-22 ---> STAT1alpha, STAT3 | 0.814 | 1.871 | 1.508 | 1.399 | 1.279 | 6.87 |
| CH000000762 | IL-22 pathway | 0.814 | 1.871 | 1.508 | 1.399 | 1.279 | 6.87 |
| CH000000579 | IL-10 ---> STAT1alpha, STAT3 | 0.814 | 1.871 | 1.508 | 1.399 | 1.279 | 6.87 |
| CH000000567 | OSM ---> ERK | 0.747 | 0.829 | 2.033 | 1.832 | 1.42 | 6.86 |
| CH000000991 | CKII ---> AP-1 | 0.991 | 1.365 | 1.065 | 2.668 | 0.765 | 6.854 |
| CH000000821 | TBK1:TRIF:IKK-i ---> p50:RelA | 1.103 | 1.675 | 1.208 | 1.52 | 1.328 | 6.835 |
| CH000004642 | TGFbetaR-I ---pak2, ERK1---> SMAD7, SERPINE1 | 1.248 | 0.623 | 2.472 | 1.292 | 1.179 | 6.814 |
| CH000004303 | NGF ---p75NTR---> p50:RelA-p65 | 0.991 | 0.866 | 1.068 | 2.884 | 1.002 | 6.811 |
| CH000000568 | OSM ---> STAT3 | 0.498 | 1.365 | 1.791 | 1.729 | 1.42 | 6.803 |
| CH000004534 | IL-6 ---> SOCS3, FOS | 0.498 | 1.365 | 1.791 | 1.729 | 1.42 | 6.803 |
| CH000004483 | IL-3 ---Ras---> Bcl-2, MBP, CREB1 | 0.807 | 0.866 | 1.644 | 1.962 | 1.487 | 6.766 |
| CH000003548 | PIAS1 ---SUMO-1---/ AR | 0.903 | 1.995 | 1.621 | 1.514 | 0.73 | 6.763 |
| CH000004690 | LXR ---/ IL1B | 1.693 | 2.2 | 1.791 | 0.986 | 0 | 6.671 |
| CH000000824 | dsRNA ---> p50:RelA | 1.075 | 1.642 | 1.171 | 1.49 | 1.279 | 6.657 |
| CH000000331 | CH000000331 | 1.693 | 0.666 | 1.791 | 1.729 | 0.765 | 6.644 |
| CH000000729 | EGF ---> STAT3 | 1.693 | 1.365 | 1.065 | 1.729 | 0.765 | 6.617 |
| CH000004525 | diacyl lipopeptide, TLR2 | 0.693 | 1.675 | 1.068 | 1.965 | 1.179 | 6.58 |
| CH000000820 | TLR3 pathway | 0.991 | 1.705 | 1.093 | 1.288 | 1.487 | 6.563 |
| CH000001010 | ITCH ---> HEF1 | 0.991 | 1.365 | 1.791 | 0.986 | 1.42 | 6.552 |
| CH000000720 | AhRR ---/ AhR | 1.187 | 1.675 | 1.291 | 1.2 | 1.179 | 6.532 |
| CH000000728 | EGF ---> STAT1alpha | 1.89 | 1.488 | 1.171 | 1.962 | 0 | 6.51 |
| CH000004595 | E-cadherin ---Eplin---> actin | 1.187 | 1.021 | 1.291 | 1.832 | 1.179 | 6.51 |
| CH000003574 | SIRT1 ---/ AR | 0.814 | 1.871 | 2.285 | 0.814 | 0.695 | 6.478 |
| CH000000026 | IL-6 ---> STAT3 | 0.589 | 1.642 | 1.334 | 1.292 | 1.586 | 6.443 |
| CH000000759 | EDAR pathway | 0.991 | 1.642 | 1.086 | 1.3 | 1.42 | 6.439 |
| CH000004536 | IL-6 signaling | 0.754 | 1.085 | 1.375 | 1.965 | 1.211 | 6.39 |
| CH000004438 | Ubc5C ubiquitin conjugation | 1.079 | 1.488 | 1.171 | 1.122 | 1.487 | 6.347 |
| CH000003545 | Chfr ---> Aurora-A | 1.079 | 1.488 | 1.171 | 1.122 | 1.487 | 6.347 |
| CH000000001 | RhoA ---/ cofilin | 1.079 | 1.488 | 1.171 | 1.122 | 1.487 | 6.347 |
| CH000000581 | deubiquitination of AF-6 | 1.079 | 1.488 | 1.171 | 1.122 | 1.487 | 6.347 |
| CH000004439 | Ubc5A ubiquitin conjugation | 1.079 | 1.488 | 1.171 | 1.122 | 1.487 | 6.347 |
| CH000000751 | T-cell antigen receptor pathway | 1.009 | 1.085 | 1.429 | 1.399 | 1.42 | 6.342 |
| CH000001027 | TopoI ---> c-Jun ---> ErbB1 | 1.234 | 1.642 | 1.334 | 1.292 | 0.837 | 6.339 |
| CH000004474 | brca1 ---> IRF7 | 1.234 | 1.642 | 1.334 | 1.292 | 0.837 | 6.339 |
| CH000000752 | Rac1 pathway | 1.079 | 1.675 | 0.952 | 0.986 | 1.644 | 6.336 |
| CH000004632 | Smad7 ---SIK---/ TGFbetaR-I | 0.814 | 1.157 | 1.508 | 0.814 | 2.039 | 6.332 |
| CH000004692 | LXR network | 1.357 | 1.871 | 2.285 | 0.814 | 0 | 6.327 |
| CH000000590 | LAT ---> p50:RelA | 0.991 | 1.559 | 1.068 | 1.383 | 1.235 | 6.235 |
| CH000004481 | IL-3 ---AKT-1---/ LNC2 | 0.628 | 1.512 | 1.502 | 1.793 | 0.792 | 6.228 |
| CH000004501 | leptin signaling | 1.327 | 1.099 | 1.171 | 1.616 | 1.012 | 6.224 |
| CH000000439 | CH000000439 | 0.903 | 1.257 | 2.472 | 1.514 | 0 | 6.146 |
| CH000004670 | oxygen independent HIF-1alpha degradation | 0.991 | 1.767 | 0.799 | 1.182 | 1.315 | 6.053 |
| CH000004726 | ID complex deubiquitylation | 1.357 | 1.157 | 0.851 | 1.399 | 1.279 | 6.044 |
| CH000004570 | ErbB1 ---Sos1, H-Ras---> Rac activation | 1.181 | 0.886 | 1.715 | 1.514 | 0.742 | 6.039 |
| CH000004494 | IFNalpha, IFNbeta ---> STAT3 | 0.689 | 1.675 | 1.291 | 1.2 | 1.179 | 6.034 |

| | | | | | | | |
|---|---|---|---|---|---|---|---|
| CH000004467 | p50:RelA-p65 ---> IL8 | 0.991 | 1.767 | 1.068 | 0.884 | 1.315 | 6.024 |
| CH000001011 | MAGI-2 ---> PTEN | 1.526 | 0.62 | 1.621 | 1.514 | 0.73 | 6.012 |
| CH000004294 | NT-3 ---trkC---> Erk | 1.013 | 0.682 | 1.596 | 1.407 | 1.279 | 5.976 |
| CH000004576 | p53 ---Ubc13---> p53{ub{K63}} ubiquitination | 1.526 | 1.257 | 0.952 | 0.877 | 1.328 | 5.94 |
| CH000000951 | CKII ---> alpha-synuclein | 0.991 | 1.365 | 1.065 | 1.729 | 0.765 | 5.915 |
| CH000004446 | TNFR1 signaling | 0.773 | 1.767 | 0.839 | 1.318 | 1.211 | 5.907 |
| CH000000772 | TNF-alpha pathway | 0.923 | 1.286 | 0.815 | 1.492 | 1.371 | 5.887 |
| CH000000878 | Src ---/ RhoA | 1.312 | 1.365 | 0.954 | 1.327 | 0.912 | 5.87 |
| CH000003888 | EGF ---> Src | 1.693 | 1.365 | 1.065 | 1.729 | 0 | 5.852 |
| CH000004175 | E1 ---UbcH7---> parkin | 0.991 | 1.365 | 1.065 | 0.986 | 1.42 | 5.826 |
| CH000004178 | E1 ---Ubc7---> parkin | 0.991 | 1.365 | 1.065 | 0.986 | 1.42 | 5.826 |
| CH000004696 | itch ---> ErbB4 ubiquitination | 0.991 | 1.365 | 1.065 | 0.986 | 1.42 | 5.826 |
| CH000000120 | CH000000120 | 0.991 | 1.365 | 1.065 | 0.986 | 1.42 | 5.826 |
| CH000003901 | Nedd4 ---> trkA | 0.991 | 1.365 | 1.065 | 0.986 | 1.42 | 5.826 |
| CH000004502 | leptin --->ERK | 0.991 | 0.666 | 1.529 | 1.383 | 1.235 | 5.803 |
| CH000000741 | Epo pathway | 1.259 | 0.767 | 1.456 | 1.292 | 0.938 | 5.712 |
| CH000000152 | CH000000152 | 1.079 | 1.488 | 2.018 | 1.122 | 0 | 5.707 |
| CH000001018 | Tip60 ---> AR | 0.589 | 1.642 | 1.334 | 1.292 | 0.837 | 5.695 |
| CH000003825 | Epicardin ---/ Rhox5 | 0.589 | 1.642 | 1.334 | 1.292 | 0.837 | 5.695 |
| CH000004188 | NRG ---> Akt-1 | 0.975 | 1.512 | 1.502 | 1.318 | 0.377 | 5.684 |
| CH000004578 | cofilin-1 degradation | 0.975 | 1.512 | 0.664 | 1.318 | 1.211 | 5.68 |
| CH000004252 | NGF ---TrkA---> Elk-1 | 0.975 | 0.651 | 1.502 | 1.318 | 1.211 | 5.657 |
| CH000000732 | EGF ---> SHIP1 | 1.079 | 1.488 | 1.171 | 1.122 | 0.792 | 5.652 |
| CH000000710 | p53 pathway | 1.234 | 1.668 | 0.613 | 1.319 | 0.815 | 5.649 |
| CH000000271 | CH000000271 | 1.234 | 1.642 | 0.613 | 1.292 | 0.837 | 5.618 |
| CH000000081 | AKT-1(h) ---> FOXO3a(h){p}:14-3-3zeta(h) | 1.234 | 1.642 | 0.613 | 1.292 | 0.837 | 5.618 |
| CH000000459 | LAP1 ---> FAK1 | 1.234 | 1.642 | 0.613 | 1.292 | 0.837 | 5.618 |
| CH000000273 | CH000000273 | 1.234 | 1.642 | 0.613 | 1.292 | 0.837 | 5.618 |
| CH000000566 | OSM ---> STAT1 | 0.45 | 1.257 | 1.621 | 1.514 | 0.73 | 5.571 |
| CH000004517 | TLR9 pathway | 0.654 | 1.157 | 1.065 | 1.292 | 1.392 | 5.561 |
| CH000000656 | insulin ---Shc---> MAPK cascade | 0.916 | 0.762 | 1.776 | 1.2 | 0.904 | 5.557 |
| CH000004479 | RIP deubiquitination | 0.814 | 1.157 | 0.851 | 1.399 | 1.279 | 5.5 |
| CH000000694 | G2/M phase (cyclin B:Cdk1) | 0.991 | 1.512 | 0.613 | 1.182 | 1.179 | 5.475 |
| CH000000537 | GEFT ---Cdc42---> c-Jun | 1.234 | 0.767 | 1.334 | 1.292 | 0.837 | 5.464 |
| CH000004540 | LC3B ---p62---/ Dvl-2 | 0.991 | 0.866 | 1.068 | 0.986 | 1.508 | 5.419 |
| CH000004720 | Endothelin-1 gene regulation | 0.498 | 0.666 | 1.791 | 0.986 | 1.42 | 5.361 |
| CH000004679 | Slim ---/ HIF-1alpha | 1.526 | 0.62 | 0.952 | 1.514 | 0.73 | 5.344 |
| CH000004482 | Bad ---> 14-3-3 | 1.526 | 1.257 | 0.952 | 0.877 | 0.73 | 5.343 |
| CH000004176 | UbcH7 ---parkin---> Eps15 | 0.903 | 1.257 | 0.952 | 0.877 | 1.328 | 5.317 |
| CH000000834 | JNK1 ---> Bax | 0.903 | 1.257 | 0.952 | 0.877 | 1.328 | 5.317 |
| CH000000958 | E1 ---/ Septin5 | 0.903 | 1.257 | 0.952 | 0.877 | 1.328 | 5.317 |
| CH000000960 | E1 ---parkin---/ synphilin-1 | 0.903 | 1.257 | 0.952 | 0.877 | 1.328 | 5.317 |
| CH000000959 | E1 ---/ synaptotagmin-11 | 0.903 | 1.257 | 0.952 | 0.877 | 1.328 | 5.317 |
| CH000000962 | E1 ---dorfin---/ synphilin-1 | 0.903 | 1.257 | 0.952 | 0.877 | 1.328 | 5.317 |
| CH000000731 | EGF ---> PI3K-C2beta | 0.903 | 1.257 | 1.621 | 1.514 | 0 | 5.294 |
| CH000000632 | PDGF A, PDGF B ---> ERK1, ERK2 | 0.958 | 0.41 | 1.531 | 1.399 | 0.946 | 5.244 |
| CH000004630 | TGFbeta1 ---> Smad2/3 | 0.814 | 0.579 | 2.285 | 0.814 | 0.695 | 5.186 |
| CH000000707 | DNA-PK ---> p53 | 1.693 | 0.666 | 1.065 | 0.986 | 0.765 | 5.175 |

| | | | | | | | |
|---|---|---|---|---|---|---|---|
| CH000004559 | CKII ---/ CARD9 | 0.991 | 1.365 | 1.065 | 0.986 | 0.765 | 5.171 |
| CH000000292 | CH000000292 | 0.991 | 1.365 | 1.065 | 0.986 | 0.765 | 5.171 |
| CH000004393 | VDR phosphorylation | 0.991 | 1.365 | 1.065 | 0.986 | 0.765 | 5.171 |
| CH000003551 | SUMO-1 ---> PIAS | 0.991 | 1.365 | 1.065 | 0.986 | 0.765 | 5.171 |
| CH000000961 | E1 ---/ ErbB3 | 0.754 | 1.085 | 1.375 | 0.725 | 1.211 | 5.15 |
| CH000004648 | angiotensin | 0.811 | 0.622 | 1.148 | 1.386 | 1.179 | 5.145 |
| CH000004656 | ERK1 ---> NQO1 | 0.991 | 0.666 | 1.065 | 0.986 | 1.42 | 5.127 |
| CH000000574 | mTOR ----> S6, eIF-4E | 0.991 | 1.488 | 1.068 | 0.986 | 0.556 | 5.088 |
| CH000000967 | PI3K ---AKT-1---/ FOXO4 | 1.079 | 1.488 | 0.558 | 1.122 | 0.792 | 5.039 |
| CH000004213 | AKT-1 ---/ FOXO3a | 1.079 | 1.488 | 0.558 | 1.122 | 0.792 | 5.039 |
| CH000004283 | ER-alpha ---CHIP---> 26S proteasome | 0.539 | 1.257 | 0.804 | 1.01 | 1.42 | 5.028 |
| CH000003412 | Akt-1 ---Mdm2---> AR | 0.589 | 1.871 | 0.613 | 1.184 | 0.765 | 5.021 |
| CH000000749 | RhoA pathway | 1.181 | 1.488 | 0.636 | 0.814 | 0.883 | 5.003 |
| CH000000662 | RhoA ---> stress fiber formation | 1.181 | 1.488 | 0.636 | 0.814 | 0.883 | 5.003 |
| CH000000889 | Fer ---> beta-catenin{Tyr142} | 0.814 | 0.579 | 1.508 | 1.399 | 0.695 | 4.995 |
| CH000000604 | Epo ---> IP3, PtsIns(3,4)P | 0.814 | 0.579 | 1.508 | 1.399 | 0.695 | 4.995 |
| CH000000947 | parkin associated pathways | 0.773 | 1.512 | 0.613 | 0.911 | 1.179 | 4.986 |
| CH000004593 | N-cadherin network | 0.991 | 0.435 | 1.068 | 1.49 | 1.002 | 4.986 |
| CH000004512 | IL-5 ---Lyn--->ERK1 | 0.903 | 0.617 | 1.369 | 0.848 | 1.179 | 4.916 |
| CH000000547 | Src ---> Grb-2 | 1.234 | 0.767 | 0.613 | 2.298 | 0 | 4.912 |
| CH000000629 | TNF-alpha ---> p50:RelA-p65 | 0.903 | 0.792 | 0.954 | 1.327 | 0.912 | 4.887 |
| CH000000769 | wnt pathway | 0.689 | 0.983 | 0.752 | 0.855 | 1.577 | 4.857 |
| CH000000121 | p105 ---> p50 | 0.573 | 1.365 | 0.613 | 1.14 | 1.105 | 4.796 |
| CH000000708 | Caspase-3 ---/ p53 | 1.526 | 0.62 | 0.952 | 0.877 | 0.73 | 4.706 |
| CH000004657 | Nrf2 ---> HMOX1 | 0.903 | 0.62 | 0.952 | 0.877 | 1.328 | 4.68 |
| CH000000822 | c-Jun degradation | 0.589 | 1.447 | 0.613 | 0.848 | 1.179 | 4.676 |
| CH000004419 | estradiol ---> E2F1 | 0.498 | 0.666 | 1.065 | 0.986 | 1.42 | 4.635 |
| CH000000740 | IFNalpha/beta pathway | 0.72 | 1.337 | 1.093 | 0.639 | 0.826 | 4.615 |
| CH000000937 | PRL ---Src,FAK1---> ERK | 1.079 | 0.543 | 0.839 | 1.414 | 0.721 | 4.596 |
| CH000004478 | IL-3 ---> c-CBL, PI3K | 0.262 | 0.836 | 1.531 | 1.399 | 0.531 | 4.56 |
| CH000000288 | CH000000288 | 1.52 | 0.836 | 0.664 | 0.634 | 0.904 | 4.557 |
| CH000003555 | NIPA ---> cyclin B1 | 0.589 | 1.393 | 0.613 | 0.814 | 1.141 | 4.551 |
| CH000000657 | insulin ---IRS---> PRK2, SHP-2, MAPK cascade | 0.655 | 0.682 | 1.596 | 0.634 | 0.806 | 4.372 |
| CH000000990 | p300 ---> ER | 0.542 | 0 | 1.171 | 1.122 | 1.487 | 4.322 |
| CH000000320 | CH000000320 | 0 | 0.767 | 1.334 | 1.292 | 0.837 | 4.23 |
| CH000000038 | insulin ---> ERK | 0.644 | 0.274 | 1.529 | 0.976 | 0.792 | 4.216 |
| CH000003940 | NMDA receptor signaling | 0.903 | 0.154 | 1.334 | 1.162 | 0.623 | 4.176 |
| CH000000864 | Htt degradation | 0.589 | 1.393 | 0.613 | 0.814 | 0.765 | 4.174 |
| CH000001028 | IGF-1 ---> Akt-1 ---> AR | 0.589 | 1.642 | 0.613 | 1.292 | 0 | 4.136 |
| CH000000760 | IGF-1 pathway | 0.589 | 1.488 | 1.068 | 0.986 | 0 | 4.131 |
| CH000000607 | BCR ---> ERK | 0.709 | 0.188 | 1.529 | 0.986 | 0.695 | 4.107 |
| CH000004628 | TGFbeta1 ---Smad7, TAK1---> p38 | 0.522 | 0.767 | 0.851 | 0.475 | 1.41 | 4.025 |
| CH000000438 | CH000000438 | 0.589 | 0.767 | 1.334 | 1.292 | 0 | 3.982 |
| CH000000994 | EGF ---> actin polymerization | 0.903 | 0.792 | 0.954 | 1.327 | 0 | 3.976 |
| CH000000250 | CH000000250 | 1.526 | 0.62 | 0.462 | 0.427 | 0.73 | 3.766 |
| CH000000249 | CH000000249 | 1.526 | 0.62 | 0.462 | 0.427 | 0.73 | 3.766 |
| CH000000032 | SP ---> Bcl-2 | 0.814 | 0 | 0.851 | 1.399 | 0.695 | 3.759 |
| CH000000262 | CH000000262 | 0.644 | 0 | 0.664 | 1.514 | 0.904 | 3.726 |

| | | | | | | | |
|---|---|---|---|---|---|---|---|
| CH000000260 | CH000000260 | 0.644 | 0 | 0.664 | 1.514 | 0.904 | 3.726 |
| CH000000461 | TGFbeta1 ---> Smad1, Smad2, Smad5 | 0.45 | 0 | 1.621 | 0.877 | 0.73 | 3.677 |
| CH000000169 | CH000000169 | 0.589 | 0 | 1.334 | 0.567 | 0.837 | 3.327 |
| CH000000151 | CH000000151 | 0.589 | 0 | 1.334 | 0.567 | 0.837 | 3.327 |
| CH000000498 | MKP-1 ---/ MBP | 0.589 | 0 | 1.334 | 1.292 | 0 | 3.215 |
| CH000004631 | SMAD7, SIK1 gene induction | 0.589 | 0 | 1.334 | 0.567 | 0 | 2.49 |
| CH000000670 | Trx1 ---> HIF-1alpha | 0.498 | 0 | 0 | 1.729 | 0 | 2.227 |

### Supplementary Table S3

**Table 3.** Medians of observed differences in pathway genes matched by pairs of patient subnetworks. The relative quantity was calculated as the number of pathway genes not shared by subnetworks divided by the number of pathway genes matched by both subnetworks combined. Calculations were performed for all pairs of patients and the 33 largest of 278 significant pathways with at least 10 pathway genes.

| Pathway | Name | Median difference |
|---|---|---|
| CH000000770 | beta-catenin network | 0.800 |
| CH000000926 | PRL pathway | 0.500 |
| CH000004648 | angiotensin | 0.500 |
| CH000000711 | TGFbeta pathway | 0.462 |
| CH000000736 | PDGF pathway | 0.429 |
| CH000000750 | insulin pathway | 0.429 |
| CH000004572 | EGF ---Raf-1---> ERK2 | 0.429 |
| CH000004475 | IL-3 signaling | 0.400 |
| CH000004506 | IL-5 pathway | 0.400 |
| CH000000772 | TNF-alpha pathway | 0.375 |
| CH000000722 | EGF pathway | 0.364 |
| CH000000710 | p53 pathway | 0.333 |
| CH000000743 | tuberin pathway | 0.333 |
| CH000000755 | TLR4 pathway | 0.333 |
| CH000000758 | stress-associated pathways | 0.333 |
| CH000000851 | JNK pathway | 0.333 |
| CH000003912 | neurotrophic signaling | 0.333 |
| CH000004149 | Nrf2 pathway | 0.333 |
| CH000000715 | HIF-1alpha pathway | 0.286 |
| CH000000751 | T-cell antigen receptor pathway | 0.286 |
| CH000000879 | Caspase network | 0.286 |
| CH000004521 | TLR2-mediated signaling | 0.286 |
| CH000000747 | E2F network | 0.273 |

| | | |
|---|---|---|
| CH000004191 | ErbB3 ---> survival | 0.273 |
| CH000000947 | parkin associated pathways | 0.250 |
| CH000001016 | AR pathway | 0.250 |
| CH000004526 | autophagy | 0.250 |
| CH000004600 | mammalian Hippo network | 0.250 |
| CH000000692 | S phase (Cdk2) | 0.200 |
| CH000000694 | G2/M phase (cyclin B:Cdk1) | 0.200 |
| CH000004525 | diacyl lipopeptide, TLR2 | 0.200 |
| CH000000752 | Rac1 pathway | 0.167 |
| CH000004341 | ER-alpha pathway | 0.143 |
